# Mammillothalamic disconnection alters hippocampo-cortical oscillatory activity and microstructure: Implications for diencephalic amnesia

**DOI:** 10.1101/590901

**Authors:** CM Dillingham, MM Milczarek, JC Perry, BE Frost, GD Parker, Y Assaf, F Sengpiel, SM O’Mara, SD Vann

## Abstract

Diencephalic amnesia can be as disruptive as the more commonly known temporal lobe amnesia, yet the precise contribution of diencephalic structures to memory processes remains elusive. We used discrete lesions of the mammillothalamic tract to model aspects of diencephalic amnesia and assessed the impact of these lesions on multiple measures of activity and plasticity within the hippocampus and retrosplenial cortex. Lesions of the mammillothalamic tract had widespread indirect effects on hippocampo-cortical oscillatory activity within both theta and gamma bands. Both within-region oscillatory activity and cross-regional synchrony were altered. The network changes were state-dependent, displaying different profiles during locomotion and paradoxical sleep. Consistent with the associations between oscillatory activity and plasticity, complementary analyses using several convergent approaches revealed microstructural changes, which appeared to reflect a suppression of learning-induced plasticity in lesioned animals. Together, these combined findings suggest a mechanism by which damage to the medial diencephalon can impact upon learning and memory processes, highlighting important role for the mammillary bodies in the co-ordination of hippocampo-cortical activity.

## Introduction

The medial diencephalon was arguably the first brain region to be linked to amnesia due to the observed pathology within this area in cases of Korsakoff syndrome (Gamper, 1928; Gudden, 1896). The importance of medial diencephalic structures for memory - in particular the mammillary bodies and anterior thalamic nuclei - has been further consolidated over the years from patient studies, non-human primate and rodent work (Dillingham et al., 2015; Jankowski et al., 2013; Vann and Nelson, 2015). However, despite the longstanding association between the medial diencephalon and amnesia, there is still no clear consensus as to why this region is so critical; given the large number of conditions in which medial diencephalic structures are known to be affected (e.g. Briess et al., 1998; Denby et al., 2009; Dzieciol et al., 2017; Kopelman, 1995; Kumar et al., 2008; Kumar et al., 2009; Ozyurt et al., 2014; Perry et al., 2019; Savastano et al., 2018; Van der Werf et al., 2000; Vann et al., 2009), this lack of knowledge presents a serious shortcoming. According to mnemonic models based on the Papez circuit, the mammillary bodies and anterior thalamic nuclei relay information from the hippocampal formation to the cingulate cortex (Delay and Brion, 1969; Papez, 1937). These models place the medial mammillary nuclei and anterior thalamic nuclei downstream from the hippocampus and, as such, attribute little intrinsic importance to these structures; instead, they are merely considered passive relays of hippocampal information (Wolff and Vann, 2019).

An alternative model is that these medial diencephalic regions are functionally upstream from the hippocampal formation, thus providing crucial inputs necessary for memory formation (Vann, 2010). While there is preliminary evidence consistent with this revised model (Vann, 2013; Vann and Albasser, 2009), it is unclear what information is provided by this ascending pathway and how it contributes to hippocampal functioning. One proposal is that the medial mammillary nuclei, by way of their inputs from Gudden’s tegmental nuclei, contribute to memory formation by setting hippocampal theta frequency (Sharp and Koester, 2008; Vann, 2009). This contrasts with earlier suggestions that the mammillary bodies simply act as a relay of hippocampal theta (Dillingham et al., 2015; Kocsis and Vertes, 1994; Tsanov et al., 2011). Alterations in hippocampal theta would be expected to co-occur with changes in cortical and hippocampal theta-gamma coupling (Canolty and Knight, 2010). Given the longstanding association between hippocampal oscillations and learning and memory related plasticity (e.g., Bikbaev and Manahan-Vaughan, 2008; Huerta and Lisman, 1993; Nokia et al., 2012; Orr et al., 2001; Tsanov and Manahan-Vaughan, 2009) an implication is that mammillary body disconnection would, in addition to altering oscillatory mechanisms, disrupt microstructural plasticity.

To test these predictions, we used a multi-faceted approach to assess the impact of mammillary body disconnection on distal areas of the Papez circuit: the hippocampus and retrosplenial cortex. Mammillothalamic tract lesions were used to model diencephalic amnesia, as damage to this tract is the most consistent feature in thalamic infarct patients with accompanying memory impairments (Carlesimo et al., 2011; Carlesimo et al., 2007; Clarke et al., 1994; Van der Werf et al., 2003; Yoneoka et al., 2004). Furthermore, this fiber tract only innervates the anterior thalamic nuclei, thus removing the possibility that any of the distal effects under investigation are simply driven by direct deafferentation of the hippocampus and retrosplenial cortex. We assessed oscillatory activity both within, and across, the hippocampus and retrosplenial cortex during theta-related states including locomotion and paradoxical sleep (Leung et al., 1982). Hippocampal dendritic spine density and clustering and the number and complexity of doublecortin-positive neurons were quantified to provide a measure of microstructural plasticity. To assess the timescale of microstructural changes, we carried out a complementary multi-timepoint magnetic resonance (MR) imaging study looking at the effects of mammillothalamic tract lesions on grey matter diffusivity. Together, these convergent approaches enable us to test current models of medial diencephalic function to determine whether they support hippocampal processes that may underpin their contribution to mnemonic processing.

## Results

### Mammillothalamic tract lesion and electrode track verification

Four separate cohorts of mammillothalamic tract (MTT) lesion rats were used (See Methods for further detail): Cohort 1 (Experiment 1/Oscillatory activity; 8 lesion, 10 controls); Cohort 2 (Experiment 2A/CA1 spines; 9 lesion, 12 controls); Cohort 3 (Experiment 2B/Doublecortin; 12 lesion, 8 controls) and Cohort 4 (Experiment 3/MR diffusivity; 13 lesion, 7 controls). Following MTT lesion, post-mortem calbindin immunoreactivity in the anteroventral thalamic nucleus is reduced (**Fig. 1F**) relative to controls (**Fig. 1E**). This, in addition to Nissl staining, provides further verification of lesion accuracy. Consistent with previous studies (Nelson and Vann, 2014; Vann, 2013), MTT lesion animals (**Fig. 1C, D**) exhibited significant spatial memory impairments both on a radial-arm maze task (Experiment 3/Cohort 4; F(1, 18) = 14.08, p = 0.001; **Fig. 1H**) and a reinforced alternation task in a T-maze (Experiment 2/Cohorts 2 & 3; supplementary **Fig. S1**).

**Figure 1.**
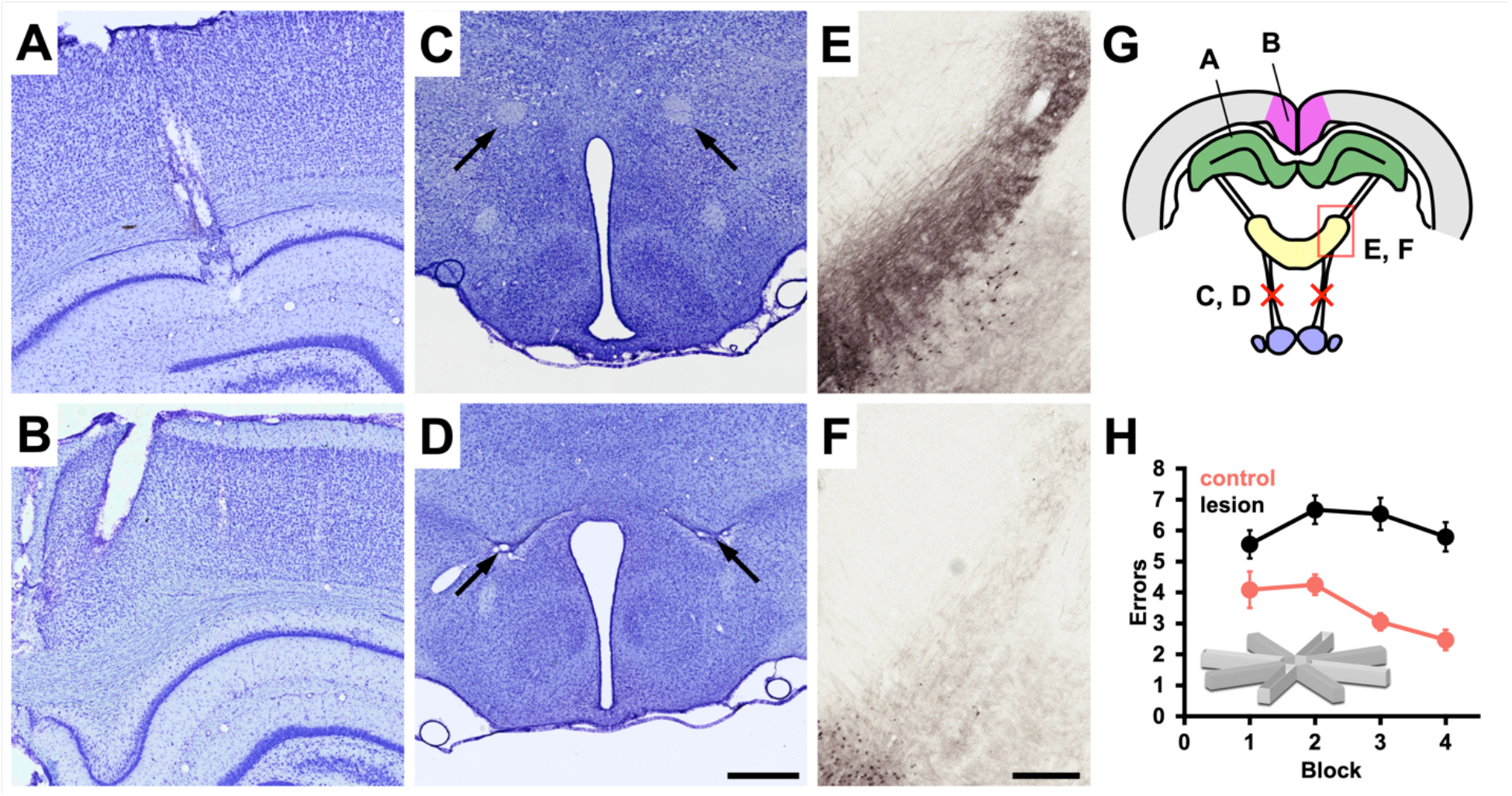
Nissl stained verification of representative electrode placements in the CA1 subfield of the hippocampus (**A**) and the retrosplenial cortex (**B**). Arrows in **B** and **C** indicate the location of the mammillothalamic tract in representative control and lesion cases, respectively, while lesion-induced reduction in calbindin immunoreactivity in the anteroventral nucleus of the thalamus (E: control – F: lesion) provided additional verification of lesion success. **G**, A schematic overview of the experimental design. **H**, Radial-arm maze (RAM) performance (from animals in the MRI cohort) demonstrates a significant lesion-induced impairment in spatial memory. Scale bar A-D: 500 µm, E-F: 250 µm.

### Experiment 1: Oscillatory Activity

To determine whether mammillothalamic disconnection influenced hippocampal and neocortical oscillatory activity, we recorded local field potentials (LFP) in the retrosplenial cortex (RSC; **Fig. 1B**) and the CA1 subfield of the hippocampal formation (HPC; **Fig. 1A**), simultaneously, in eleven rats (supplementary **Fig. 1**). Six of these animals had received discrete bilateral lesions of the mammillothalamic tract (arrows; Fig. 1C, D). An additional seven animals, two of which had mammillothalamic tract lesions, were implanted in the RSC alone. The remaining five surgical controls were recorded singularly from either brain region (HPC, n = 2; RSC, n = 3).

#### Mammillothalamic tract lesions attenuate hippocampal theta frequency during locomotion

To promote locomotion across an evenly distributed, wide range of running speeds, animals were trained to retrieve sugar pellets in a bow-tie maze, which resulted in animals consistently reaching speeds of 40-55 cm/s. No significant difference was found between lesion and control groups in the density distributions of speed measurements (supplementary Fig. S6; 0-55 cm/s; p = 0.47). Theta power, and frequency, are both positively correlated with running speed in HPC (Carpenter et al., 2017; Jeewajee et al., 2008; Maurer et al., 2005; Sheremet et al., 2016; Slawinska and Kasicki, 1998). To determine whether disconnection of the mammillary bodies from the Papez circuit impaired the encoding of speed in HPC and RSC, we first examined the power and peak frequency of theta oscillations from recordings in awake, locomoting animals. As expected, the peak theta-band frequency of control animals in both the HPC (0.03 Hz ± 0.00, t = 10.95, p < 0.001; **Fig. 2B, C**) and RSC (0.03 Hz ± 0.00, t = 11.42, p < 0.001; **Fig. 2E, F**), showed highly significant positive linear correlations with running speed. By contrast, the peak theta frequency of lesioned animals was significantly attenuated across all running speed bins in both the HPC (−0.57 Hz ± 0.15, t = −3.79, p = 0.003; **Fig. 2B, C**), and the RSC (−0.59 Hz ± 0.16, t = −3.63, p = 0.003; **Fig. 2E, F**). Mammillothalamic lesions attenuated the peak frequency of theta but they did not disrupt the relationship between running speed and theta frequency (treatment/speed interaction, RSC: −0.00 ± 0.00, t = −1.71, χ = 2.92, p = 0.087; HPC: −0.00 ± 0.00, χ = 0.33, t = −0.58, p = 0.583; **Fig. 2B, E**), reflecting, at least in part, intact glutamatergic/GABAergic septo-hippocampal connections (Carpenter et al., 2017; Fuhrmann et al., 2015).

**Figure 2.**
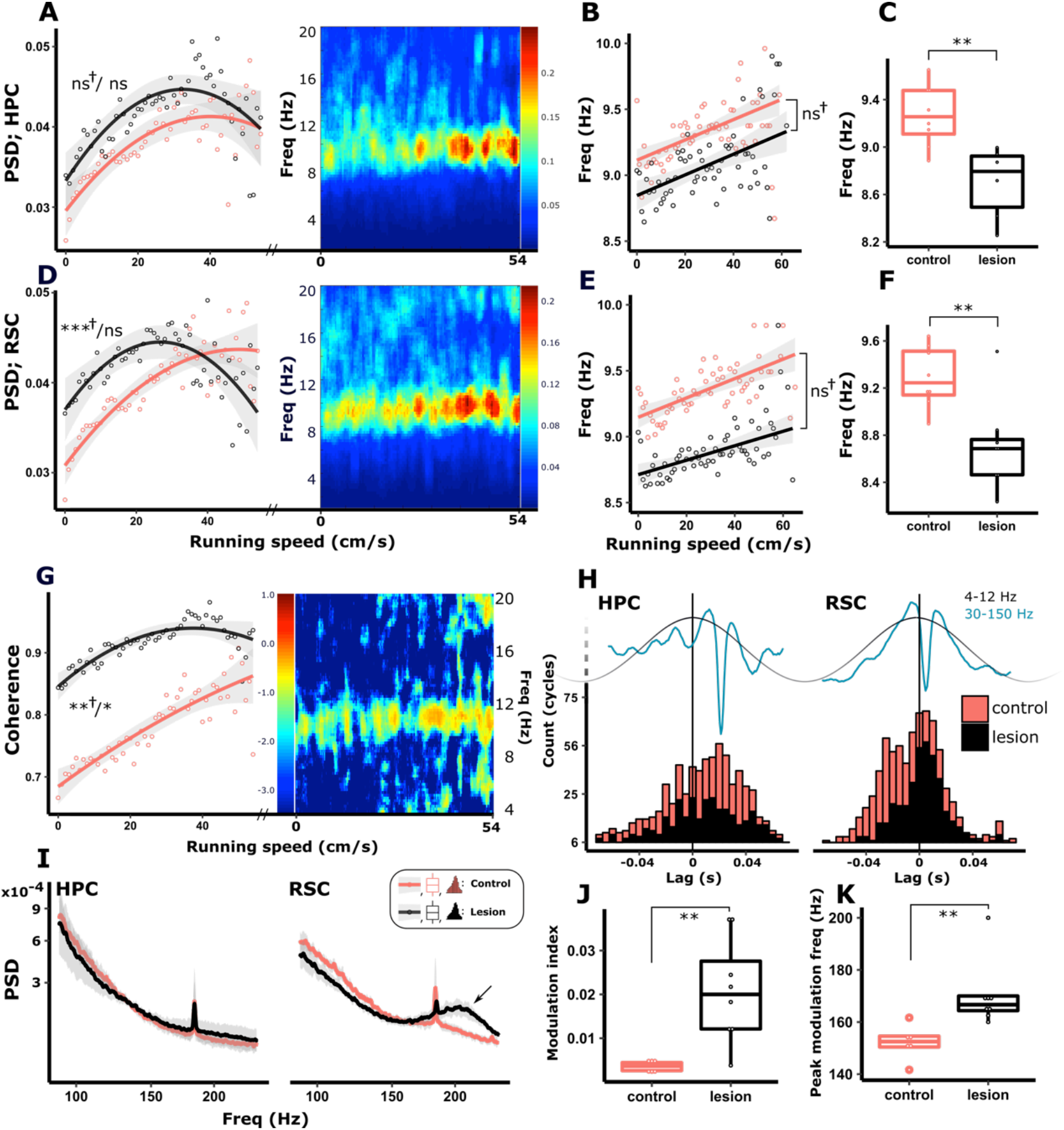
In control animals, both peak theta frequency and theta power (PSD) are modulated by running speed in hippocampus (HPC; **A**-**C**) and retrosplenial cortex (RSC; **D**-**F**). In animals with MTT lesions, HPC power-speed relationship was intact (**A**). Although the relationship between peak theta frequency and running speed was intact in lesioned animals (**B**, HPC; **E**, RSC), peak theta frequency was significantly attenuated across the range of measured speeds in both HPC (**C**) and RSC (**F**). Theta-band coherence between HPC and RSC is positively correlated with running speed (**G**). Heat maps in **A**, **D** and **G** show higher theta power in HPC, RSC, and theta-band coherence between the two regions, respectively, increasing with speed in individual, representative control animals. Theta-band coherence between HPC and RSC in lesioned animals was significantly greater across speed bins (**G**), however animals with lesions had an intact relationship between theta-coherence and running speed (**G**). (**H**) Gamma band (30-150 Hz) trough-triggered averages of theta (4-12 Hz) oscillations were used to calculate the time lag from the theta cycle peak. Consistent with previous studies (Belluscio et al., 2012), gamma power was most strongly associated with the descending phase in HPC and the peak of the theta cycle in RSC with no difference between treatment groups. Gamma power during locomotion was unaffected by MTT lesions, however increased power in the 180-240 Hz range (high frequency oscillations; HFO) was observed in RSC (arrow in the right facet of **I**). Corresponding increases in phase amplitude coupling in the corresponding frequency range were observed in RSC (**J**, **K**), reflecting the increased HFO. Abbreviations: **, p<0.01; *, p<0.05; ns, not significant; † alongside significance represents results of speed/treatment interactions.

While running speed and theta frequency exhibited a linear relationship, that of theta power and running speed was best represented by a quadratic function (**Fig. 2A, D, G**), likely reflecting the plateau in interneuron- and pyramidal cell firing rate at higher running speeds (i.e. > 20 cm/s; Ahmed and Mehta, 2012; Maurer et al., 2005). Theta power in both HPC (2.30×10^−4^ ± 1.47×10^−5^, t = 15.64, χ = 187.81, p < 0.001; **Fig. 2A**) and RSC (2.08×10^−4^ ± 1.48×10^−5^, t = 14.04, χ = 157.24, p < 0.001; **Fig. 2D**) were significantly correlated with running speed and no overall difference between theta-band power was found in either region between treatment groups. In HPC, no difference was observed in the relationship between power and running speed (**Fig. 2A**) but in RSC, the speed/power relationship was significantly reduced in magnitude (−2.66×10^−6^ ± 1.21×10^−6^, t = −2.21, χ = 45.17, p < 0.001; **Fig. 2D**). Theta-band coherence, between HPC and RSC, increases with running speed and, consistent with the relationship between theta power and running speed, plateaus at higher speeds (>30 cm/s). Absolute coherence between RSC and HPC in animals with MTT lesions was significantly increased when compared across all speed bins (0.15 ± 0.05, t = 2.99, p = 0.017) and, in addition, the relationship between coherence and speed was attenuated in rats with MTT lesions (−5.46×10^−5^ ± 1.85×10^−5^, t = −2.96, χ = 9.24, p = 0.010; **Fig. 2D**; supplementary Fig. S5).

During locomotion, gamma power and frequency (Ahmed and Mehta, 2012) as well as coupling between theta phase and gamma amplitude (Sheremet et al., 2019) are positively correlated with running speed. Hippocampal gamma is thought to reflect the local interaction between inhibition and excitation generated from the interaction between pyramidal neurons and interneurons. Considering the effects that we had established in the theta frequency band, we next examined gamma and higher frequency oscillations (HFO; >100 Hz). Looking at coupling between the phase of theta and the amplitude of gamma oscillations we first established that, consistent with results of Belluscio et al. (2012) and Koike et al. (2017), gamma oscillations are consistently dominant on the descending slope of the theta cycle while RSC gamma is dominant closer to the theta peak (**Fig. 2H**). Using the time points of the troughs of gamma oscillations (>2 std. deviations of the baseline power) to generate the average of broadband filtered (0-500 Hz) LFP, we measured the time lag between theta peak and gamma trough (**Fig. 2H**) and found no difference between treatment groups. While temporal coupling was maintained in both regions, both the modulation index, derived from the phase amplitude coupling, as well as the gamma/high frequency activity of modulation, was significantly increased in the RSC of lesioned animals (**Fig. 2I-K**). As expected, this finding was the result of an increase in high frequency power centered at ∼170 Hz. (**Fig. 2K**; see also supplementary **Fig. S7**). It is worthy of note that such high frequency oscillations (i.e. >120 Hz) may not reflect true oscillatory activity but instead may be, in part, influenced by multiunit activity around the electrode tip (Merker, 2013; Ray and Maunsell, 2011; Scheffer-Teixeira et al., 2013). Therefore, in as much as these effects may be classified as inhibitory/excitatory-related oscillatory imbalance, they may equally be classified as local high frequency hyperactivity.

#### Mammillothalamic tract lesions reduce Running Speed-Related Theta Asymmetry

To better understand the nature of the observed attenuation in theta frequency in MTT animals, we next looked at the dynamics of the theta cycle. Hippocampal theta cycles have an inverse saw-toothed shape with a shorter duration ascending phase (trough-to-peak) and a longer descending phase (peak-to-trough), resulting in an asymmetric waveform (Buzsaki et al., 1985) (**Fig. 3A, B)**. The asymmetry index (absolute difference in the duration of ascending and descending phases) within HPC theta oscillations was found to be positively correlated with running speed (0.46 ± 0.25 ms, t = 18.53, χ = 241.24, p < 0.001; **Fig. 3C** upper facet), resulting principally from a decrease in ascending phase duration (−0.25 ± 0.13 ms, t = −19.73, p < 0.001; **Fig. 3E**). Consistent with our finding that MTT animals showed an intact speed/peak theta frequency relationship (**Fig. 2B**), HPC theta ascending phases in lesioned animals were also significantly correlated with speed, however, ascending phase durations across all speed bins were significantly longer (6.72 ± 2.53 ms, t = 2.66, p = 0.021; **Fig. 3C, E**; supplementary **Fig. S3**).

**Figure 3.**
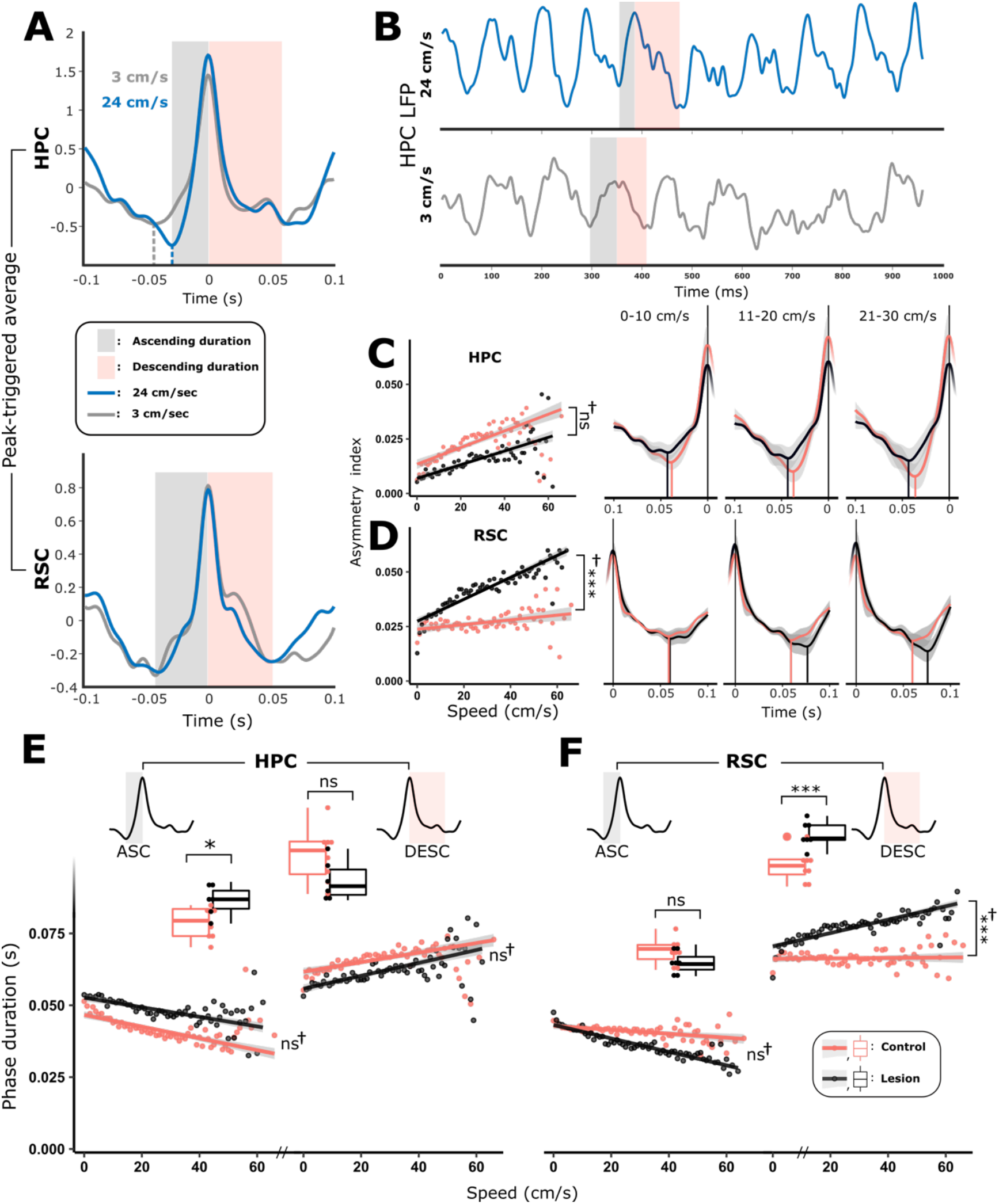
The hippocampal (HPC) theta cycle is an asymmetric, sawtooth waveform with a short ascending phase, and a longer duration descending phase (**B**). Cycle asymmetry becomes exaggerated with increased rates of locomotion (**A**-**C**). The absolute asymmetry (asymmetry index; AI) in the HPC theta cycle is highly correlated with running speed (**D**), reflecting an attenuation in ascending phase duration (**E**, **F**) and an increase in descending phase duration (**E**). Consistent with their observed peak frequency attenuation, MTT animals exhibited longer duration ascending phases than controls (**E**, left facet). Control RSC ascending phases decrease in duration with increasing speed (**F**) with consistent duration descending phases. In the RSC of MTT animals, cycle asymmetry is significantly more positively correlated with speed than in controls, resulting primarily from an increase in descending phase duration (**C**, lower facets; **F**)

In control animals, the strength of correlation between asymmetry index and running speed was much weaker in RSC than in HPC, but it was statistically significant (0.12 ± 0.027 ms, t = 4.35, χ = 18.46, p < 0.001). However, we found that in lesioned animals, the RSC asymmetry index showed a considerably stronger increase with speed compared to control animals (0.37 ± 0.034 ms, t = 10.64, χ = 105.41, p < 0.001; **Fig. 3D**). This mainly arose from a greater increase in descending phase duration with speed (0.23 ± 0.02 ms, t = 10.13, χ = 96.18, p < 0.001; **Fig. 3F**) and a greater attenuation in ascending phase duration in the lesion animals (−0.16 ± 0.02 ms, t = −8.70, χ = 72.15, p < 0.001). Phase entrainment of RSC interneurons is independent of hippocampal input (Talk et al., 2004). Recent modelling of the dynamics of spike-phase coupling in the context of theta cycle asymmetry (Cole and Voytek, 2018) present an interesting case that the duration of phase within a cycle is related to the firing frequency of phase-locked neurons such that longer duration descending phases in RSC may provide for a longer window in which firing activity may reach peak frequency, thus resulting in increased frequency activity. In the context of our findings, such a mechanism may serve to link the lesion-related changes in descending phase duration in RSC (**Fig. 3F**; right facet) with the observed increase in 180-240 Hz HFO (**Fig 2I**; right facet).

#### Mammillothalamic tract lesions attenuate theta frequency during paradoxical sleep

Next, to establish whether the lesion-related group effects on theta during locomotion were consistent across other high-theta states, LFP recordings were made in the HPC and the RSC during paradoxical (REM) sleep (PS). During periods of PS, MTT lesion rats did not differ from controls in HPC theta power, however, a significant attenuation in theta frequency was apparent in lesioned animals (−0.32 Hz ± 0.11, t = −2.96, p = 0.013; **Fig. 4A, B, C**). In the RSC, lesioned animals had a significantly higher theta power but with no discernible difference in peak theta frequency (**Fig. 4D, E, F**). Theta cycles during PS were more symmetrical than during awake locomotion (Fig. 4G; also see supplementary **Fig. S7**) and no difference between lesion and control groups was found in either the asymmetry index or ascending/descending phase durations in either HPC (**Fig. 4H**), or RSC (**Fig. 4I**).

**Figure 4.**
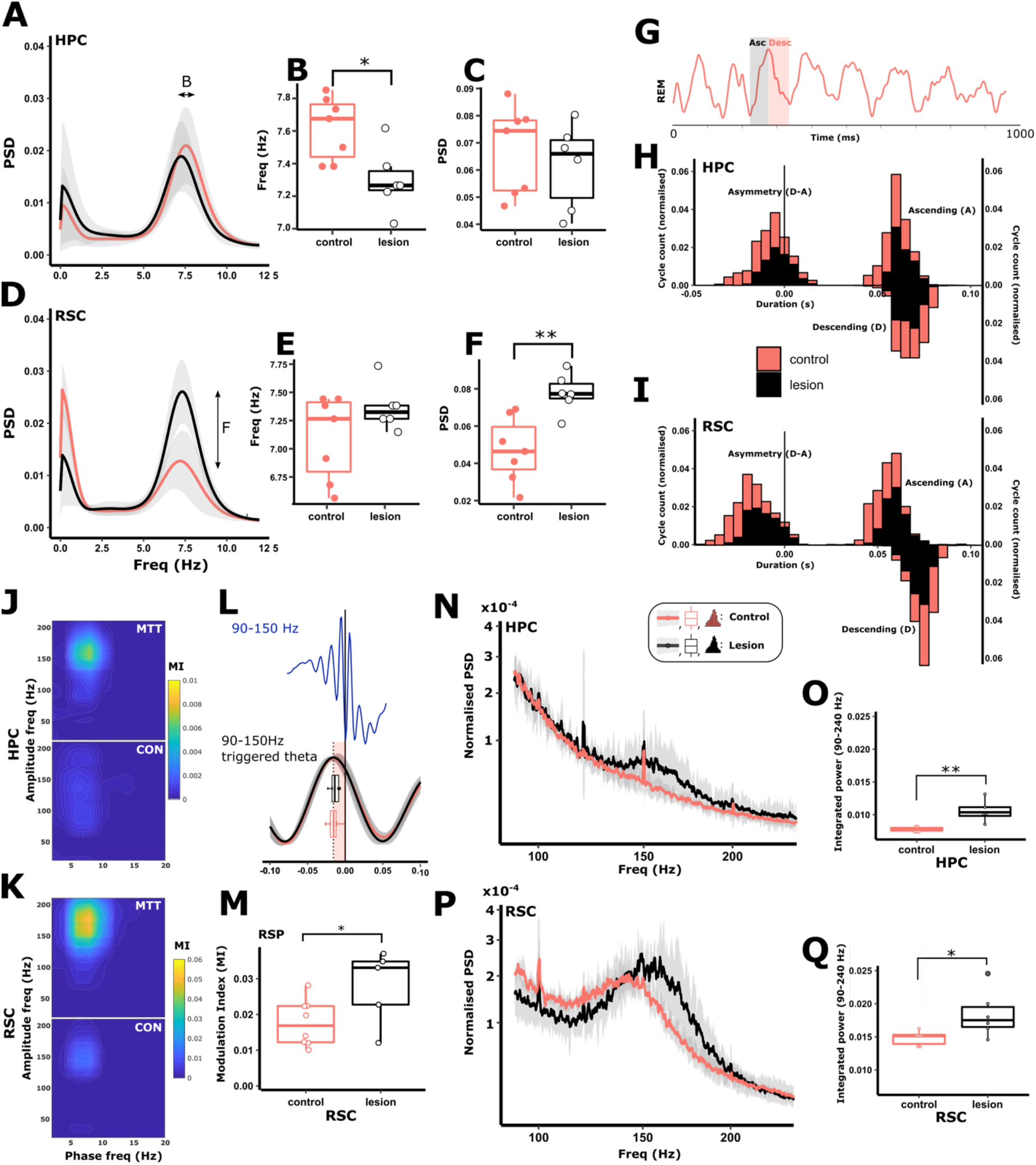
During paradoxical sleep (PS) hippocampal theta frequency, but not power, was significantly attenuated (**A**, **B, C**), while cortical theta power, but not frequency, was significantly greater (**D, E, F**). **G** shows a typical LFP trace from HPC during PS (shaded grey, ascending phase; shaded red, descending phase). **H**, **I**; Lesion of the MTT did not influence the absolute asymmetry of theta cycles in either the HPC (**H**; left histogram) or the RSC (**I**; left histogram) as reflected by consistent, overlapping distributions of ascending and descending phase durations in both HPC and RSC (right histograms in **H** and **I**, respectively). During PS, strong theta-gamma phase-amplitude coupling is present in the fast gamma frequency range (90-150 Hz; **J**). The degree to which theta and fast gamma are coupled (modulation index; MI) was significantly increased in both HPC (**J**) and RSC in lesioned animals (**K**), while the time lag between the peak of the theta cycles, and the troughs of fast-gamma oscillations (>2 standard deviations of the baseline power) remained consistent between treatment groups (**L**). **L** shows the average theta waveform triggered by the trough of fast-gamma (90-150 Hz) oscillations (time = 0 s). Shaded red area highlights the time lag between the trough of the gamma oscillations and the peak of the preceding theta peak with corresponding box plots showing the very similar means, and variance of time lags compared between lesion (black) and control (red) animals. Lesion-dependent increases in phase-amplitude coupling in both HPC (**N**, **O**) and RSC (**P**, **Q**) were, therefore, the product of increased high frequency power at frequencies that corresponded to the coupling observed in **J** and **K**.

During PS, RSC theta phase is strongly coupled to the amplitude of fast gamma/HFO (80-150 Hz) amplitude (Koike et al., 2017; Fig. 4J), with weaker phase amplitude coupling in HPC. The modulation index (MI) is a quantification of the strength of coupling between theta phase and the amplitude of gamma/HFO (Tort et al., 2010). Consistent with these findings, during PS, control animals showed strong theta-gamma/HFO phase-amplitude coupling during PS in RSC and a weaker MI in HPC (0.01 ± 0.00, t = 2.45, p = 0.044; **Fig. 4J**) and RSC (0.01 ± 0.00, t = 2.23, p = 0.048; **Fig. 4K, M**).

In the RSC, the amplitude of gamma oscillations/HFO (30-150 Hz) is strongly coupled with the descending phase of the theta cycle during PS (Belluscio et al., 2012; Koike et al., 2017). First, to determine whether the phase (within the theta cycle) in which fast gamma was dominant was affected by disconnection of the mammillary bodies, we calculated the time lag between the trough of gamma oscillations (of power > 2 standard deviations of the baseline), and the peak of the synchronous theta cycle. No significant group difference was observed in the mean lag time (**Fig. 4L**), suggesting that theta-entrainment of cortical unit activity was unaffected by MTT lesions; this is in line with the finding that RSC entrainment to theta is independent of HPC input (Talk et al., 2004). The increased phase-amplitude coupling between theta and fast gamma was, therefore, likely due to higher fast gamma power in MTT animals. To test this, we looked at the power spectral density in the frequency band in which phase-amplitude coupling peaked (90-240 Hz). Within this fast-gamma band, overall gamma power across this band was significantly higher in both HPC (0.003 ± 0.001, t = 3.621, p = 0.007; **Fig. 4N, O**) and RSC (0.004 ± 0.002, t = 2.40, p = 0.037; **Fig. 4P, Q**) of MTT animals.

### Experiment 2: Hippocampal spine and doublecortin analysis

#### Mammillothalamic tract lesions reduce the number and clustering of CA1 spines and the number and complexity of doublecortin-positive neurons in dentate gyrus

Since oscillatory activity and learning-induced plasticity are tightly coupled (e.g. Orr et al., 2001; Tsanov and Manahan-Vaughan, 2009) we hypothesized that changes in local field potential might reflect altered structural plasticity in the hippocampus. To address this possibility, we examined dendritic spine density and clustering in Golgi-stained tissue in one group of rats (Cohort 2) and the expression of doublecortin (DCX), a marker of adult neurogenesis (Couillard-Despres et al., 2005), in a second group following training on a T-maze task (Dudchenko, 2001) (Cohort 3).

MTT lesions reduced the density of spines on basal CA1 segments (t(19) = 2.23, p = 0.038; **Fig. 5A**). Furthermore, the distribution of CA1 spine clustering was significantly attenuated in lesioned compared with control animals (t(19) = 2.37, p = 0.028; **Fig. 5B**). In contrast to the changes in CA1, there were no changes between lesion groups in either the density (t(19) = 0.38, p = 0.71), or the clustered distribution (Mann-Whitney U = 42.00, p = 0.39) of spines in the apical segments of RSC pyramidal neurons (supplementary **Fig. S8B, C**). Both spine density and clustering are associated with learning and memory in intact animals (Harland et al., 2014; Moser et al., 1994; Moser et al., 1997); spine clustering is thought to underlie stronger and more efficient information processing in dendrites (Kastellakis et al., 2015; Rogerson et al., 2014). The reduction in spine density and clustering in the lesion animals may, therefore, be an additional contributor to impoverished encoding in this lesion model.

**Figure 5.**
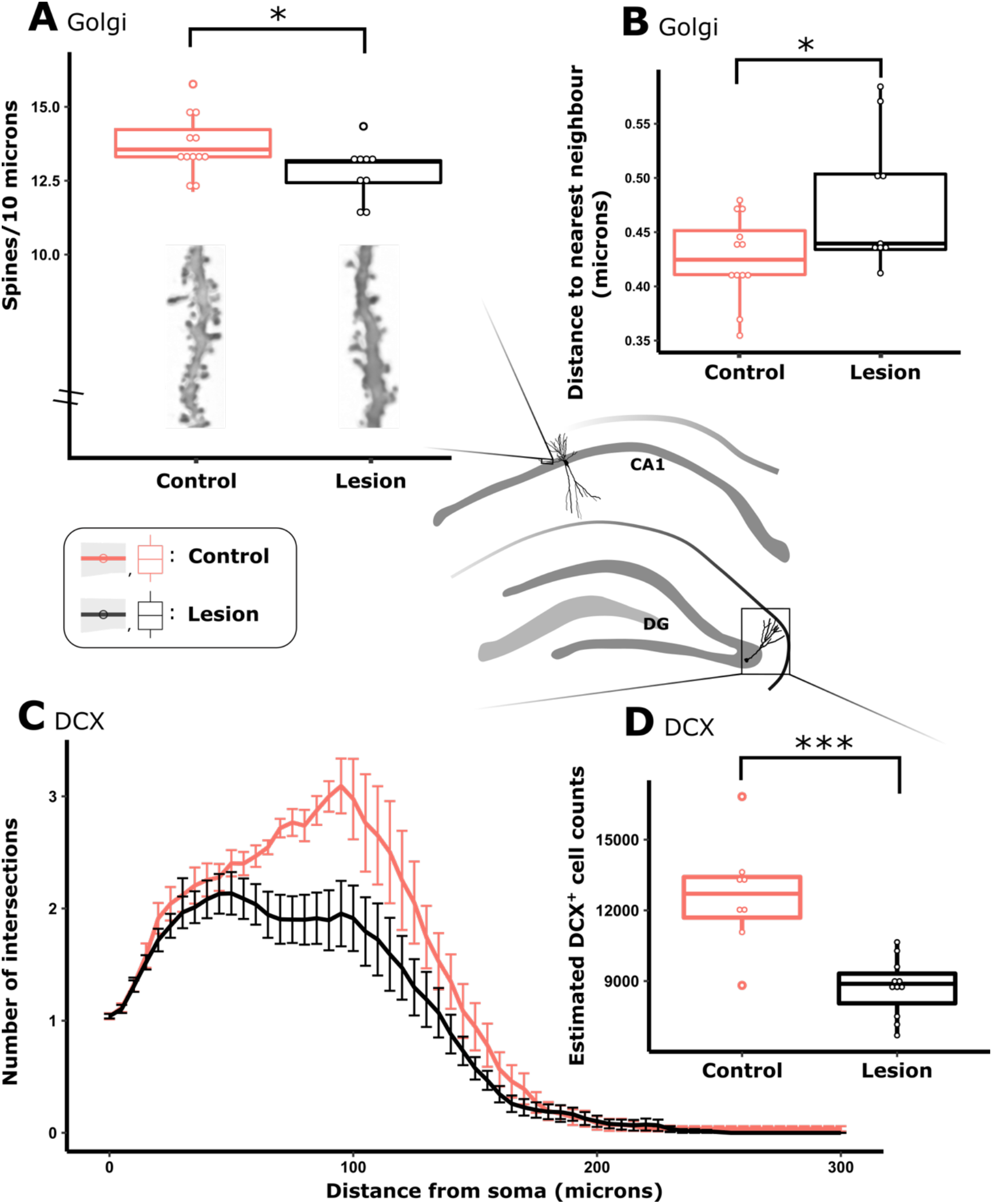
Mammillothalamic tract lesions were associated with a decrease in both number (**A**) and clustering (**B**) of spines in the basal dendrites of Golgi stained CA1 pyramidal neurons. Doublecortin (DCX) is a marker of adult neurogenesis; in a separate cohort of animals, lesions resulted in a decrease in the complexity of DCX^+^ neurons (**C**) as well as a decrease in the stereologically-estimated overall number of DCX^+^ neurons in the dentate gyrus (DG; **D**). Central schematic diagram depicts the morphology and anatomical location of neurons that were sampled.

Spatial training has been linked to structural changes within existing hippocampal cells but also to increased numbers and dendritic arbor complexity of adult-born neurons in the dentate gyrus (Kempermann et al., 2004; Vukovic et al., 2013). MTT lesions reduced the unbiased stereological estimation of the number of dentate gyrus DCX cells by approximately 31% (t(17) = 4.76, p < 0.001, **Fig. 5D**). In addition to there being fewer DCX cells in the lesion group (**Fig. 5D**), those cells that were present, exhibited less morphological complexity, as reflected by a reduced number of intersections (Sholl analyses; F(1, 18) = 9.05, p = 0.008; Radius main effect (p < 0.001); interaction p = 0.07; **Fig. 5C**). Given that no difference was observed in the complexity of Golgi-stained dentate gyrus neurons (see supplementary **Fig. S8A**), this effect is likely to be limited to newly formed cells. The reduction in DCX cells may be a contributing factor to the memory impairments associated with MTT lesion as targeted experimental suppression of hippocampal neurogenesis impairs performance on spatial memory tasks (Dupret et al., 2008; Farioli-Vecchioli et al., 2008; Lemaire et al., 2012).

### Experiment 3: MR diffusivity

#### Diffusion-weighted imaging and microstructural plastic changes

The reductions in doublecortin-positive cells and spine density were observed following training on a spatial memory task. We therefore hypothesized that MTT lesions led to the attenuation of experience-driven structural plasticity and tested this prediction by employing a longitudinal magnetic resonance imaging (MRI) study. Spatial learning has been found to evoke changes in grey matter diffusivity in the hippocampal formation of both humans and rodents. Moreover, in one study, altered diffusivity co-occurred with enhanced staining for various plasticity markers such as synaptic boutons, glial reactivity and BDNF levels (Sagi et al., 2012; Tavor et al., 2013). Diffusion-weighted MRI therefore offers a non-invasive means of investigating correlates of experience-driven plastic changes in intact and disease model animals.

#### Brain-wide lesion-induced diffusivity changes

Rats (Cohort 4) were scanned at four separate time-points: prior to surgery (Scan 1); nine weeks following MTT-lesion surgery or control surgery (Scan 2); at the beginning of radial-arm maze (RAM) training (Scan 3); at completion of RAM training when the controls were proficient at the task (Scan 4; **Fig. 6C**). We employed a voxel-wise mixed ANOVA to uncover brain-wide interactions between group (lesion vs control) and scan time (four levels) for two complementary diffusivity metrics, fractional anisotropy (FA) and mean diffusivity (MD) (Feldman et al., 2010). Since differential changes in MD (supplementary **Fig. S9**) appeared to derive mainly from the ventricular system (presumably a direct effect of ventricular enlargement following the lesion), this measure was not analyzed further. On the other hand, voxels where differences in FA values reached significance (p < 0.05) were found across many large white matter structures such as the corpus callosum, cingulum, fornix and the mammillothalamic tract (**Fig. 6A**). Clusters of significant voxels were also present in grey matter including the hippocampus, mammillary bodies, posterior granular retrosplenial cortex and parts of the midbrain.

**Figure 6.**
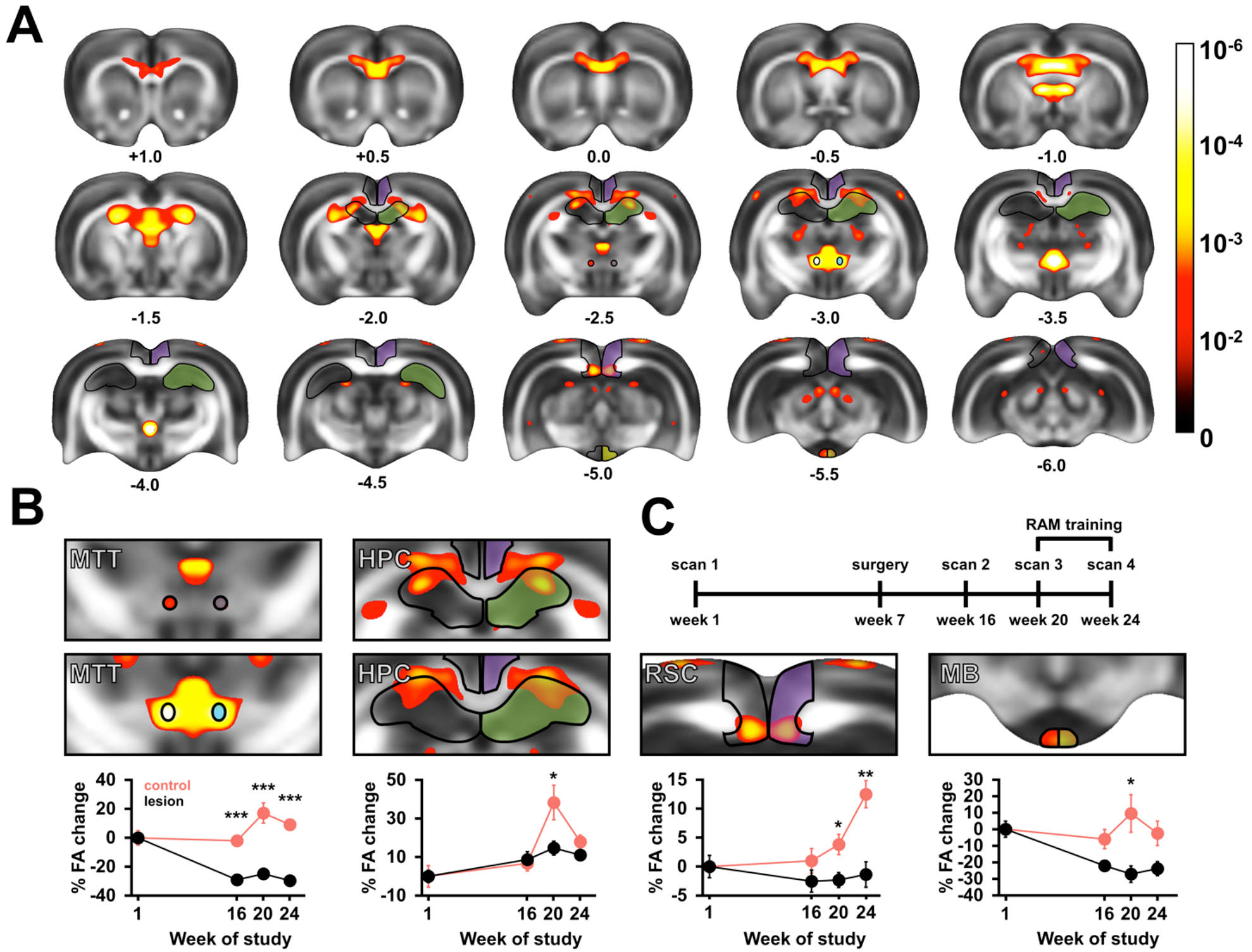
Changes in fractional anisotropy (FA) elicited by MTT lesion and radial-arm maze training. (**A**) Results of an ANOVA with the between factor of surgery (two levels) and the within factor of scan number (four levels). Significant voxels have been thresholded to α < 0.01 and displayed using a heat map overlaid on top of a mean FA template. Successive coronal levels are numbered according to distance from bregma (mm). Outlines designate anatomical structures from which regions of interest for regional analyses were selected (at α < 0.005). The calibration bar represents p-values on a logarithmic scale (−Log_10_) (**B**) The timeline of regional changes in the mammillothalamic tract (MTT), hippocampus (HPC), retrosplenial cortex (RSC) and the mammillary bodies (MB). The y-axis displays change from mean baseline (prospective lesion and sham cases) values at Scan 1; error bars represent SEM. (**C**) Timeline of scanning sessions. Animals were scanned at four timepoints over a six-month period; surgeries were carried out during week 7.

#### Higher hippocampal and retrosplenial FA following spatial learning in intact animals but not in lesion cases

We next performed regional post-hoc comparisons to examine the timeline of diffusivity changes more closely. The selected regions of interest comprised clusters at α < 0.005 (uncorrected, see Methods) present in the anterior dorsal hippocampus (HPC), posterior granular retrosplenial cortex (RSC), posterolateral medial mammillary bodies (MB) and, for comparison, in the mammillothalamic tract (MTT; **Fig. 6B**).

Consistent with previous reports, spatial training elicited higher FA in intact animals (Sagi et al., 2012). While the MTT and HPC values peaked following initial training (MTT: p = 0.035, HPC: p < 0.001), the RSC values showed a gradual increase, reaching maximum levels following the final test session (p = 0.008 at Scan 3 and p < 0.001 at Scan 4). This complementary pattern of diffusivity changes is consistent with the medial diencephalon and hippocampus supporting memory encoding (Vann and Aggleton, 2003) with the retrosplenial cortex having a greater role in long-term memory storage (Milczarek et al., 2018). By contrast, lesion cases exhibited markedly attenuated FA in relation to initial training (MTT: p < 0.001, MB: p < 0.043, HPC: p = 0.011, RSC: p = 0.018) as well as following the final training session in the RSC (p = 0.001). While the first post-surgery scan (Scan 2; pre-training) revealed altered FA values in the MB and MTT (regions that have sustained primary and secondary damage; p < 0.001) the differences in HPC and RSC, sites distal to the lesion, were not observed until animals had undergone spatial training. We therefore interpret these distal changes in diffusivity as the failure of lesion cases to express task-evoked plasticity.

## Discussion

The traditional view of the Papez circuit is that the mammillary bodies and anterior thalamic nuclei are functionally downstream of the hippocampus, acting principally as a hippocampal relay (Aggleton and Brown, 1999; Barbizet, 1963; Papez, 1937). Instead, the present results support a revised model whereby the mammillary bodies have an important role in optimizing hippocampal-cortical oscillations, in particular modulating the frequency of theta, the level of inter-regional coherence and, in turn, local hippocampal and cortical activity. Concordantly, MTT lesions diminish a number of markers of learning-induced plasticity, which together likely contribute to the marked memory impairments observed.

Discrete mammillothalamic tract (MTT) lesions were used to model diencephalic amnesia, enabling us to disentangle the contribution of the mammillary bodies without concomitant damage to the supramammillary nuclei. This is important given the influence of the supramammillary nuclei on hippocampal theta frequency (e.g., Kirk, 1998). As such, many studies that have assessed the effects of mammillary body lesions, or inactivation, on hippocampal theta can be difficult to interpret when damage extends into the overlying supramammillary nuclei (Sharp and Koester, 2008; Zakowski et al., 2017) and vice versa (Renouard et al., 2015). Nevertheless, the only previous study that has examined hippocampal theta following mammillary body manipulation in non-anesthetized animals established a similar pattern of changes to those reported here, with a reduction in hippocampal theta-frequency (Sharp and Koester, 2008). The lesions in Sharp and Koester’s study included both the mammillary bodies and supramammillary nuclei, however, our present findings would suggest their results were at least in part driven by the loss of the mammillary bodies. Theta frequency is considered a mechanism for timing the flow of information in the hippocampus (Richard et al., 2013) and high-frequency theta in particular has been linked to spatial memory in both rodents and humans (Goyal et al., 2018; Olvera-Cortes et al., 2002). Furthermore, reducing the frequency of theta in otherwise normal animals impairs performance on spatial memory tasks (Pan and McNaughton, 1997). The implication, therefore, is that this reduction in hippocampal theta frequency would contribute to the memory impairment observed in MTT lesion animals.

In our revised model of the mammillary body contribution to the Papez circuit, the medial mammillary nuclei integrate theta input from the ventral tegmental nuclei of Gudden. Thus, to more fully characterize any dysfunction associated with disconnection of this theta stream, we assessed hippocampal-cortical activity across two theta-rich states: awake locomotion, and paradoxical sleep (PS). During locomotion, several characteristics of theta (e.g. frequency, power, cycle waveform characteristics) are correlated with running speed and, in the most part, these associations were spared by mammillary body disconnection, e.g. theta frequency was attenuated overall but its relationship with running speed was preserved.

Not only was there a frequency attenuation in the lesion animals during locomotion but the hippocampal theta cycle in these animals exhibited a more symmetrical waveform, characterized by longer-duration ascending (trough-to-peak) phases. As in HPC, RSC theta asymmetry is positively correlated with running speed (albeit to a lesser degree than in HPC). An unexpected finding of the study was that the RSC descending phase duration of lesioned animals increased dramatically with running speed relative to controls (**Fig. 4F**). This change was accompanied by a large increase in HFO (180-240 Hz) power and significantly higher phase-amplitude coupling at a frequency corresponding to the spectral peak of the HFO. Inhibitory interneurons fire preferentially in the descending phase of the theta cycle (Buzsaki, 2002). Extrapolating from the findings of Cole and Voytek in the HPC (2018), this pattern of effects may be explained by relating the RSC hyperactivity to the observed changes in the dynamics of the theta cycle such that with longer descending phases, phase-locked interneurons have an increased ‘preferred-phase window’ within which to reach their intrinsic firing frequency, resulting in a greater maximally attained firing frequency across each cycle.

In control animals, hippocampal theta cycles during PS are more symmetrical than during locomotion (supplementary **7A-C**). As a result, longer ascending phase durations of awake locomoting lesioned animals were found to be closer to control PS cycles than those of control locomoting animals (supplementary **Fig. 7A**), suggesting that in lesions animals, theta cycles are less able to vary between states. Silencing septal theta during PS dramatically attenuates hippocampal theta power and disrupts contextual memory (Boyce et al., 2016). While mammillary body disconnection did not disrupt hippocampal theta to the same degree (Boyce et al., 2016), it did lead to an attenuation in theta frequency during PS (supplementary **Fig. 3D**), potentially highlighting a previously unexplored involvement of these structures in consolidation. While HPC theta frequency was reduced in the lesion animals both during locomotion and PS, the RSC showed some state-dependent changes in theta. In contrast to the reduced theta frequency during locomotion, the frequency of PS theta in the RSC was not reduced and, instead, a significant increase in theta power was observed (**Fig. 4D, F**). The increased HFO in the RSC observed during locomotion in MTT lesion animals was also found during PS (supplementary **Fig. 3G**). Given the peak frequencies of HFO during PS was considerably lower than during locomotion, these changes are likely to reflect the induced hyperactivity of separate subpopulations of the RSC that are specifically active during either PS (68% of RSC units reported by Koike et al., 2017) or during locomotion. Unlike in awake animals, however, no effect was seen in waveform dynamics of the theta cycle, leaving open an explanation for the PS-associated hyperactivity (in both HPC (Fig. 4N, O) and RSC (supplementary Fig. S7). One mechanism by which mammillary body projections could influence the frequency of hippocampo-cortical theta frequency is through the cholinergic system. Following chemogenetic inhibition of cholinergic neurons in the medial septum, Carpenter and colleagues (2017) reported an attenuation in theta frequency in the hippocampal formation that mirrored the magnitude and direction of that which we report here following lesions of the mammillothalamic tract. Both hippocampal and cortical cholinergic activity is reduced following mammillary body lesions (Beracochea et al., 1995) and while the mammillothalamic projection itself is not cholinergic, it may act to modulate cholinergic activity through interactions either with its direct connections with the anteroventral thalamic nucleus/nucleus basalis of Meynert, or through indirect influence on distal brain regions. One such pathway, described by Savage and colleagues (2011) in explanation of reduced cholinergic activity in HPC following lesions of the anterior thalamus, is via the modulation of descending RSC projections to the medial septum (Gonzalo-Ruiz and Morte, 2000).

Lesion induced hyperactivity in RSC, particularly during PS, may reflect a more synchronized, i.e. deactivated, cortical state in which there is a reduction in the variance of population-wide activity (Harris and Thiele, 2011). Both the observed increase in theta power during PS, the corresponding increase in HFO, as well as conserved theta phase coupling support a case for increased local synchronicity. Animals with MTT lesions also exhibited excessive inter-regional coherence across both awake and PS states, which again likely reflects a reduction in functional diversity in the system following the loss of ascending mammillary body projections. Indeed such excessive synchrony is thought to reduce information coding capacity (Cagnan et al., 2015) while increased regional connectivity has been observed in neurological conditions linked to impaired cognitive ability (Gardini et al., 2015; Hawellek et al., 2011).

Several distal microstructural differences were also observed in MTT lesioned rats, with fewer spines present on the basal dendrites of CA1 pyramidal neurons, and a corresponding reduction in spine clustering (see **Fig. 5**); the latter a mechanism that, in normal animals, likely serves to strengthen neural responses (Kastellakis et al., 2015). Rats with MTT lesions had significantly fewer doublecortin-positive neurons in the dentate gyrus, and those that were present showed a reduced morphological complexity, suggesting that mammillothalamic disconnection may be associated with a reduction in adult hippocampal neurogenesis (Boyce et al., 2016). Given that all of these measures increase in normal animals following spatial training (Lemaire et al., 2012; Moser et al., 1997; Tronel et al., 2010; Vukovic et al., 2013), this combination of observed microstructural differences in lesion animals may well reflect a reduction in the capacity for learning-induced plasticity. Such an explanation is further supported by our diffusion MR study where the distal impact of lesioning the MTT became evident only once animals underwent spatial training, suggesting the loss of learning-induced plasticity in lesion cases. Together, these convergent findings reinforce the pervasive effects of MTT lesions on numerous aspects of neural plasticity.

There is increasing evidence of mammillo-thalamic pathology in neurological conditions that present with memory impairments (e.g. Denby et al., 2009; Dzieciol et al., 2017; Kumar et al., 2008; Kumar et al., 2009; Perry et al., 2019) reinforcing the need to better understand the contributions of these medial diencephalic structures. Together, our results provide insight into mechanisms via which the mammillary bodies can contribute to memory processes: ascending mammillary projections optimize hippocampo-cortical plasticity via their contribution to oscillatory architecture. Furthermore, given the changes observed during PS, it is possible that this region is not only involved in encoding, as previously assumed (Vann and Nelson, 2015), but may also have additional roles in memory consolidation. If distal oscillatory disturbances underlie the memory impairments following medial diencephalic damage, it raises the possibility that artificially restoring the electrophysiological patterns to normal could serve as a therapeutic intervention. Deep brain stimulation of the hypothalamus, fornix and medial temporal lobe has been reported to improve memory function and increase plasticity and neurogenesis (Lee et al., 2013); the current findings, therefore, provide a possible mechanism via which these memory improvements may occur.

## Methods

### Animal subjects

#### Experiment 1 – Oscillatory activity

Twenty-five naïve male Lister-Hooded rats (Envigo, UK) were used weighing 300–350 g at the time of surgery (Cohort 1). Of these animals, 8 were completely excluded from this study due to either misplaced electrodes (n = 3), incomplete MTT lesions (n = 1), or technical issues resulting from attempts to co-house post-surgery (n = 3). Of the remaining 17, 8 received complete bilateral MTT lesions, electrode implantation in CA1 and RSC, as well as chronic placement of electromyography (EMG) electrodes. Five surgical controls received the same electrode configuration without MTT lesions, an additional 4 received electrode implantation in CA1 (n = 2) or RSC (n = 2) but were also implanted in the claustrum for participation in a separate study (see supplementary **Fig. S6**).

#### Experiment 2 – Hippocampal spine and doublecortin analyses

For the Golgi spine analyses (Cohort 2), Naïve male Lister-Hooded rats were used (Harlan, UK). Rats weighed 230 – 300 g at the time of surgery, 14 received bilateral MTT lesions and 12 underwent control surgery. Five animals had sparing of the MTT and were removed from all subsequent analyses. For the doublecortin analyses (Cohort 3), Dark Agouti rats (Harlan, UK) were used weighing 191 – 225 grams at the time of surgery, 16 received bilateral MTT lesions and 8 underwent control surgery. Four of the 16 rats had bilateral sparing and were removed from subsequent analyses. One lesion case was excluded from the doublecortin cell number estimates analysis due to large air bubbles obscuring part of the dentate gyrus on a number of sections.

#### Experiment 3 – MR diffusivity

Twenty-six naïve male Lister-Hooded rats (Envigo, UK) were used weighing 230 – 280 g at the time of surgery (Cohort 4). Sixteen animals received bilateral MTT lesions and 10 underwent control surgery. Three animals had sparing of the MTT and were removed from all subsequent analyses; two control animals died during the study and MR data from one control animal could not be recovered.

The total number of rats used in this study was 101 with a final N of 78 after rats with sparing of the MTT, misplaced electrodes, or other technical issues were excluded. Following surgery, the electrophysiology rats were housed singly while all other rats were housed in small groups of 2 – 4 rats per cage under diurnal light conditions (14 h light/10 h dark) with free access to water and environmental enrichment. During behavioral testing, animals were food deprived to within 85% of their free feeding weight. Animal husbandry and experimental procedures were carried out in accordance with the European Community directive, 86/609/EC, and the Cruelty to Animals Act, 1876, and were approved by the Comparative Medicine/Bioresources Ethics Committee, Trinity College, Dublin, Ireland, and followed LAST Ireland and international guidelines of good practice (**Experiment 1**). **Experiments 2-3** were carried out in accordance with the UK Animals (Scientific Procedures) Act, 1986 and associated guidelines, the EU directive 2010/63/EU, as well as the Cardiff University Biological Standards Committee.

### Surgical procedures

Surgeries were performed under an isoflurane-oxygen mixture (induction 5%, maintenance 2 – 2.5% isoflurane). Once anesthetized, animals were positioned in a stereotaxic frame (David Kopf instruments, California, US).

#### Mammillothalamic lesions

With bregma and lambda level, the scalp was incised to expose the skull. Bilateral craniotomies were then made at positions targeting the mammillothalamic tract (AP: bregma −2.5 mm, ML: bregma ±0.9 mm DV: −6.9 mm (from top of the cortex). A thermocouple radiofrequency electrode (0.7-mm tip length, 0.25-mm diameter; Diros Technology Inc., Canada) was lowered into position and radiofrequency lesions were made (70°C/33 seconds using an OWL Universal RF System URF-3AP lesion maker; Diros Technology Inc., Canada). For sham surgeries, the probe was lowered to 1.0 mm above the target in order to avoid damaging the tract; no lesion was made.

#### Electrode implantation

If the surgery involved electrode implantation, the skull was cleaned before 5-6 screws were secured and cemented to the skull. Craniotomies were made before careful removal of the dura and subsequent implantation of electrodes in positions corresponding to the following coordinates (mm from bregma unless stated): (CA1: AP-3.6: ML: ±3.4; DV: −1.9 from top of cortex with a 18.5° angle towards the midline; RSC: AP: −3.6; ML: ±1.1; DV: −1.2 from top of cortex with a 24° angle towards the midline). Rats were then implanted with combinations of 6 or 7 tetrode bundles (25 µm platinum–iridium wires; California FineWire, CA, USA) mounted onto a drivable 32-channel microdrive (Axona Ltd., St. Albans; UK). An additional 1-2 × custom made twisted bipolar electrodes (70 µm stainless steel), were incorporated into the remaining microdrive channels via 34 AWG, PTFE-insulated wire (Omnetics, MN, USA). The same wire was adapted for electromyographic (EMG) recordings from beneath the nuchal muscle. Implanted microdrives and bipolar electrodes were secured in place using dental cement (Simplex Rapid; Dental Medical Ireland; Dublin, Ireland).

The scalp was then sutured, and antibiotic powder was topically applied (Acramide: Dales Pharmaceuticals, UK). Animals were rehydrated with a subcutaneous 5-10 ml injection of 4% glucose saline, and given post-operative analgesia (0.06ml, 5 mg/ml Meloxicam, Boehringer Ingelheim, Rhein, Germany).

### Behavioral experiments

#### T-maze

Prior to testing, all animals in Experiment 2 were given 2 days of habituation to the T-maze. Animals were then given either 10 (Cohort 2) or 6 (Cohort 3) sessions of training. One session was conducted each day, with eight trials in each session. Each test trial consisted of a forced ‘sample’ phase followed by a ‘choice’ phase. During the forced sample phase, one of the goal arms of the T-maze was blocked. After the rat turned into the pre-selected goal arm, it received a reward. Animals were then returned to the start arm for 10 s. During the choice phase animals were given free choice to enter either arm, only receiving a reward if the direction opposite to the forced choice in the sample run was chosen (i.e., non-matching to sample choice). Left/right allocations for the sample runs were pseudo-randomized over daily trials, sessions and rats, with no more than three consecutive sample runs to the same side in each session. The start arm was kept constant for the whole procedure.

#### Radial-arm maze

Animals first received 4 habitation sessions which involved unrestricted exploration of all arms of the maze, each containing a scattering of sucrose pellets (45 mg; Noyes Purified Rodent Diet, U.K.). Rats were trained on a working memory version of the radial-arm task that involved the retrieval of sucrose pellets from each of the arms of the maze. At the start of the trial, all arms were baited, and the animal was placed on the center platform and allowed to make an arm choice. After returning to the center platform, all doors were shut for 10s before being re-opened and the animal could then make another arm choice. This procedure continued until all arms had been visited or 10 min has elapsed. Rats received two trials/session every other day, totaling 32 trials over 16 sessions.

### Perfusion/histology

At the end of behavioral experiments, animals were given an overdose of sodium pentobarbital (60 mg⁄kg, Euthatal, Rhone Merieux, Harlow, UK) and transcardially perfused with 0.1 M phosphate buffer saline (PBS) followed by 4% paraformaldehyde (PFA) in 0.1 M PBS. Brains from all experiments other than Experiment 2A were removed and post-fixed in 4% PFA for 4 h before being transferred to 25% sucrose in distilled water overnight. On the following day, 40µm sections were cut in the coronal plane using a freezing-stage microtome. One 1-in-4 series was collected directly onto gelatin-subbed slides and Nissl stained (Cresyl Violet, Sigma-Aldrich, Gillingham, UK) for verification of lesion location. One series was subsequently processed for doublecortin staining, a marker for young neurons. The remaining series were reacted for doublecortin (rabbit polyclonal, Abcam, UK; 1:1000 dilution) and calbindin (D28k; mouse monoclonal; 1:10,000; Swant).

#### Golgi staining

Golgi staining of the whole brain was performed according to the manufacturer’s instructions (FD Rapid GolgiStain Kit™; FD NeuroTechnologies, Columbia, MD). Mounted 150 µm coronal sections cut on a cryostat were counterstained with Cresyl Violet, dehydrated in an ascending series of alcohols, cleared in xylene, and coverslipped using DPX mounting medium (Thermo Fisher Scientific, UK). Slides were re-coded by a researcher not involved in data collection to allow blinded analyses.

#### Immunohistochemistry

Sections were washed 4×10 min in 0.1 M PBS containing 0.2% Triton X-100 (PBST) between each incubation period. Endogenous peroxidase activity was quenched in a 0.3% hydrogen peroxide solution (0.3% H_2_O_2_, 10% methanol, and distilled water) for 5 min. Sections were then incubated in M PBS containing 3% normal serum for 1 hour followed by a 48 hour primary antibody incubation at 4°C with 1% normal serum. Following 3×10 min PBST washes, sections were reacted for 2 hours in a 1:250 secondary solution containing 1% normal serum. After a further 3×10min PBST washes, sections were reacted for 1 hour in an avidin/biotin-peroxidase complex solution with 1% normal serum (ABC Elite, Vector Laboratories, Peterborough, UK). Sections were then washed twice in 0.05M Tris buffer and the label was visualized with 3,3-diaminobenzidine kit (DAB Substrate Kit, Vector Laboratories, Peterborough, UK) according to the supplier’s protocol. Sections were then mounted on gelatin coated slides, dehydrated in an ascending series of alcohols, cleared in xylene and coverslipped with DPX mounting medium.

## Data analysis

### Experiment 1: Electrophysiology

Recordings were performed during the light phase of a 12 hour, day/night cycle. Following a recovery period of no less than 8-10 days rats were trained to retrieve sugar pellets from wells in four end compartments of a bow-tie maze to promote locomotion over a broad range of speeds. Once rewards were retrieved from one end of the maze, a manually operated middle door was opened allowing the animal to investigate and retrieve rewards from the opposite end. Local field potential recordings were obtained using the Axona DacqUSB acquisition system (Axona Ltd, St. Albans, UK) at a sampling frequency of 4.8 KHz and down-sampled to 960 Hz and standardized for all further analyses. Animal positional information obtained using a light bar mounted with infra-red LEDs, one larger than the other, was sampled at 250 Hz using a ceiling mounted infra-red video camera. Positional information was interpolated to match the sampling frequency of the down-sampled LFP. Both prior to- and following recordings in the bow-tie maze, rats were socialized with litter-mates for an average of 1-2 hours to provide natural sleep deprivation following which, recordings were performed on rats in a familiar square home-arena where they could sleep.

All analyses were performed using custom scripts in MATLAB (v. 2018b; MathWorks).

#### Power spectral density (PSD)/coherence

Welch’s method (*pwelch* function) was used for calculation of frequency domain spectral power and the coherence between signals was estimated using magnitude-squared coherence (*mscohere* function).

#### Running speed analysis

Running speed was calculated based on the change in position over time calculated from the animal’s positional information (sampled at 200 Hz). Running speed was first resized through cubic interpolation to match the sampling frequency of the LFP (960 Hz). The average running speed within 600 ms windows of LFP was calculated for individual recording sessions. Windows were then sorted and assigned speed values based on nearest-integer rounding. Window bins of common speed were concatenated, and speed bins of less than 2 seconds combined duration were discarded. Five, 1.8 second windows of each speed bin were randomly extracted for subsequent analyses.

#### Theta cycle asymmetry

Determination of the phase of a signal using bandpass filtering and the Hilbert transformation, for instance, temporally homogenizes the oscillatory activity, masking cycle asymmetry (Belluscio et al., 2012; Buzsaki et al., 1985). To preserve and measure oscillatory asymmetry, theta cycle windows were identified first through identification of zero crossing time points in LFP signal that had been bandpass filtered between 4-12 Hz. Zero crossings time points were then used to extract start and end points for positive (0-180°) and negative oscillation phases (180-360°), which were in turn used to extract windows of the same LFP to which a broader bandpass filter had been applied (0.5-80 Hz). Within positive and negative amplitude windows, the time points of maximum and minimum amplitudes, respectively, were extracted and rescaled to give peak and trough times from which ascending (trough-to-peak) and descending (peak-to-trough) durations were calculated. The absolute difference in ascending and descending durations provided an asymmetry index.

#### Paradoxical sleep detection

Paradoxical sleep (PS) is characterized by high theta power, low delta power as well as muscle atonia. Episodes of PS were indexed by first identifying awake periods in which rats were moving (running speed>0). A moving-window (12 sec with 50% overlap) based multitaper method (Mitra and Bokil, 2008) was used on bandpass-filtered (1-60 Hz) LFP for time-domain calculation of spectral power. A threshold based on the theta(4-12Hz))-delta(1-4Hz) power ratio (TD ratio) was then calculated and epochs of SWS and PS were characterized by a low TD ratio (high delta, low theta power), and high TD ratio, respectively. Episodes of putative PS (high TD ratio; minimum 15 seconds in duration) were validated using a threshold set for the root mean square of the synchronously recorded EMG signal (see supplementary **Fig. S4**).

#### Phase-amplitude coupling

Phase-amplitude coupling was calculated using the approach of Tort and colleagues (2010). Briefly, theta phase (2° bins ranging from 0-20°) was calculated through the Hilbert transformation, creating a homogenized theta cycle from which gamma/high-frequency oscillations (HFO) ranging from 30-200° (in 5° bins), was compared. An adaptation of the Kullback-Leibler distance was used to generate a modulation index (MI), derived from a phase-amplitude plot.

### Experiment 2: Golgi and doublecortin analysis

#### Golgi Sholl analysis (Cohort 2)

Image stacks were obtained with a Zeiss LSM 510 confocal microscope (Carl Zeiss Ltd, UK) using a 20x apochromat objective. A Helium-Neon laser (633 nm) was used to image the slices at high resolution (1024 × 1024 pixels). Images were acquired at pre-set intervals (0.5 µm) on the Z-plane, so generating image stacks (about 120-140 images per stack) that allowed the analysis of dendritic arbors in three-dimensions. ImageJ (1.48a Fiji version, NIH, Bethesda, MD)(Schindelin et al., 2012) and its free segmentation plugin Simple Neurite Tracer (Longair et al., 2011) were used for semi-automated tracing and Sholl analysis of dendritic arbors. The analysis involved overlaying the cell with a series of concentric spheres spaced 1µm apart, centered on the cell body, and counting the number of dendritic arbors intersecting each circle relative to the distance from the soma to a maximum distance of 300 µm for Rgb neurons and 350 µm for CA1 and DG neurons.

Eligible arbors were required to be intact, clearly visible, clear of artefacts, and well-isolated from neighboring neurons. In Rgb, Sholl analyses were performed on the dendritic arbors of small fusiform and canonical pyramidal neurons whose somata were positioned in superficial layers II-III of the rostral portion of Rgb while in CA1, basal arbors of CA1 pyramidal neurons were used. In the DG Sholl, analyses were performed on neurons whose somata were located in the granule cell layer and whose dendritic arbors were projecting towards the molecular layer.

Dendritic arbors were traced from a total of 476 cells, 262 from surgical control brains and 214 from MTT-lesioned brains with at least five arbors for each of apical Rgb, basal Rgb, basal CA1, and DG per animal. For Rgb arbors, identifying cells with both apical and basal segments that were eligible for tracing was rare (seven from the MTT-lesioned group and one from the control group) due to staining artefacts or overlapping of soma or dendrites from neighboring neurons. Specifically, 73 basal Rgb arbors and 77 apical Rgb arbors for the Sham group, and 46 basal Rgb arbors and 54 apical Rgb arbors for the MTT-lesioned group were traced. For hippocampal arbors, 59 CA1 and 53 DG arbors for the Sham group, and 65 CA1 and 49 DG arbors for the MTT-lesioned group were traced.

#### Dendritic spine counts (Cohort 2)

Image stacks were acquired with a Leica DM6000 microscope with a digital camera (Leica, DFC350 FX) and Leica Application Suite imaging software. A 100x (NA 1.4) oil-immersion objective employing transmitted light was used to collect 30 – 90 images per stack with a µm step-size. Dendritic segments of between 20 µm and 25 µm were traced and cropped with the Simple Neurite Tracer plugin (Longair et al., 2011) according to the eligibility criteria adapted from Harland et al. (2014): one segment per neuron was counted; segments did not belong to the primary branch; segments were unobscured by other dendrites or staining artefacts; segments started and ended equidistant between two spines and started at least 10 µm from any terminal or branching points. Stacks of images were then processed using a custom ImageJ (Fiji v1.51h, https://imagej.net/) macro. Briefly, stacks were filtered and sharpened and collapsed into a single 2D projection plane for manual counting of spines. Cartesian coordinates of identified spines were transformed to map onto a linear representation of the dendritic branch and spine density (number of spines/10 um dendrite length) and mean nearest neighbor (distance to the nearest spine in 1-dimensional space) were calculated. In total 249 Rgb apical and 347 CA1 basal segments were included (4579 Rgb spines; 9242 CA1 spines).

#### Doublecortin Sholl analysis (Cohort 3)

Only mature (late phase) cells (Mandyam et al., 2008) were analyzed; the inclusion criteria were as described above. Plane images were captured under bright-field microscopy with a Leica DMRB microscope equipped with a 20x (NA 0.50) objective, a CCD camera (Olympus, DP73) and Cell Sense Dimensions software 1.16 (Olympus). Images were converted to 8-bit greyscale and inverted before Sholl analyses (to a maximum distance of 300 µm from the soma). Dendritic arbors were traced from a total of 158 cells, 88 cells for MTT lesioned cases and 70 cells from surgical controls with 5-16 cells analyzed per case.

#### Doublecortin cell number estimates (Cohort 3)

The unilateral total number of doublecortin neurons was estimated using the optical fractionator method (Gundersen, 1986; West et al., 1991), performed using a Leica DM6000 microscope equipped with a motorized stage (BioPrecision, Ludl Electronic Products), Z-axis focus control (Ludl Electronic Products, #99A420), and a CCD camera (CX 9000) connected to a computer running StereoInvestigator 8.0 software (both MicroBrightField, Williston, VT). Sampling of sections followed a systematic, uniform random sampling scheme. A section sampling fraction of 1/4 was employed resulting in 15 – 23 sections sampled per brain (cut section thickness 40µm; mounted section thickness 13 µm). The contour of the GCL including the subgranular zone covering the entire dentate gyrus extending 1.80 – 6.60 mm posterior to bregma (Paxinos and Watson, 1998) was traced live using a 10x objective (0.4 NA) and the cells were counted using a 63x oil immersion objective (NA 1.4). A dissector height of 8µm with a 2µm guard zone was used to sample in the z-axis. The counting frame area was 2500µm^2^ (50 x 50µm) and the x – y step length was set to 100µm (area 10,000µm^2^). These parameters resulted in a mean of 347 total doublecortin neurons (range, 266 – 447) being counted in an average of 639 counting frames (range, 471 – 817). The number of doublecortin neurons was estimated from.

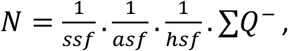

where *ssf* is the section sampling fraction; *asf*, the area sampling fraction, is the ratio of the counting frame area to the area of the x – y step length; *hsf*, the height sampling fraction, is the ratio of the dissector probe height to the Q^−^ weighted tissue thickness; and ∑ Q^−^ is the total count of cells sampled (Dorph-Petersen et al., 2001).

The coefficient of error (CE) was calculated (Gundersen et al., 1999) to determine whether the sampling method was sufficiently precise. As a measure of the variability of the estimates the observed inter-individual coefficient of variation (CV = standard deviation/mean) was determined. The sampling strategy was considered optimal as the biological variation contributed more than 50% to the observed relative variance, thus providing a reliable indication of the true numbers of DCX cells: 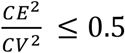 (see **Table S1**).

### Experiment 3: MR Diffusivity

*Scans:* Animals received 4 MR scanning sessions: 6 weeks before surgery (Scan 1); 9 weeks after surgery Scan 2); following the second session of radial-arm testing (13 weeks from surgery; Scan 3); and following the final session of radial-arm testing (17 weeks from surgery; Scan 4). Prior to scanning and behavioral testing, rats were placed in a quiet, dark room for 60, and 30 min, respectively.

Rats were scanned with a 9.4 Tesla MRI machine (Bruker, Germany) with a 72 mm, 500 W four-channel transmit coil. The protocol comprised a multi-gradient structural T2 RARE scan (8 repetitions; 24 slices at 500 µm z-spacing and an xy resolution of 137 µm; duration: 10 min 40 s), followed by a DTI scan with a diffusion-weighted spin-echo echo-planar-imaging (EPI) pulse sequence. The DTI acquisition parameters were TR/TE=4000/23.38 ms, 4 EPI segments, 32 gradient directions with a single b-value at 1000 s/mm2 and five images with a b-value of 0 s/mm2 (B0) (Jones et al., 1999). Each scan included 24, 500 µm coronal brain slices of with an x-y resolution of 273 µm. The EPI acquisition lasted 40 min.

During scan acquisition, rats were lightly anesthetized with 1-1.5% isoflurane in medical air and placed on a heatmat. The heart rate, breathing rate and temperature of the animals was monitored throughout each scanning session. The entire scanning protocol lasted just over an hour and DTI images were acquired within 80-100 min of the end point of behavioral testing.

#### Extraction of DTI metrics and normalization procedures

Diffusion weighted data were processed through a combination of custom Matlab (Mathworks, USA) scripts, and the *ExploreDTI* toolbox (Leemans et al., 2009). Briefly, this comprised a motion/distortion correction by co-registration of diffusion weighted volumes to the initial B0 image (with appropriate b-matrix rotation (Leemans and Jones, 2009), correction for Gibbs ringing artefacts (Osher and Shen, 2000; Sarra, 2006), and robust tensor fitting (Chang et al., 2005). Subsequently, metrics of fractional anisotropy (FA) and mean diffusivity (MD) (Pierpaoli and Basser, 1996) were extracted.

Image normalization was achieved using the Advanced Normalization Tools (ANTS) software package (Avants et al., 2011). A cohort specific template was first created through iterative co-registration of FA maps. Co-registration of individual FA volumes to the template was then achieved through symmetric diffeomorphic registration (Avants et al., 2008) with reuse of the resultant warp fields to provide equivalent normalization of MD data. In all cases a cubic b-spline interpolation scheme was used.

#### Creation of mean hemispheric maps

Inspection of the generated FA and MD maps revealed asymmetric noise related to the acquisition procedure. To remedy this, mean hemispheric maps were produced (e.g. Kikinis et al., 2019). This approach was deemed appropriate due to, a) the bilateral lesioning of the ipsilaterally-projecting mammillothalamic tract (Vann & Albasser, 2009) symmetrical to the midline and, b) the assumption that lesion-induced and training-induced changes would not show laterality (e.g. Sagi et al., 2012). The maps were further processed in ImageJ (Fiji v1.51h, https://imagej.net/). First, the images were resized to a higher resolution (factor of three) by employing bicubic interpolation. Next, coronal slices were flipped horizontally and pairs of mirror images were iteratively registered to their mean to increase the hemispheric symmetry (rigid registration, followed by affine registration and elastic transformations (Sorzano et al., 2005) (https://imagej.net/BUnwarpJ)). The co-registered mirror images were mean-averaged and a mean FA template was created. The template was then manually masked to restrict the analyses to brain tissue only. All FA and MD maps were finally smoothed using a 0.3 mm 3D Gaussian kernel.

#### Statistical analyses

Statistical analyses on data obtained in Experiment 1 were carried out in R Statistics (version 3.5.2; provided in the public domain by the R Foundation for Statistical Computing, Vienna, Austria; R Development Core Team, 2009, available at http://www.r-project.org/)). General linear models (“lme4” package; Bates and Maechler, 2010) were fitted for lesion/control group comparisons. For comparisons involving speed/treatment interactions, a random term was included to account for repeated measures. Given the difficulties associated with determining degrees of freedom in these instances, p-values were obtained through likelihood ratio tests comparing nested models, i.e. inclusion of the individual components of the interaction but not the interaction itself. All graphs were generated using the “ggplot2” package (Wickham and Chang, 2012; http://cran.R-project.org/web/packages/ggplot2), while figures were compiled in Inkscape (Inkscape v.0.92.4, The Inkscape Project, freely available from www.inkscape.org).

For Sholl analyses the number of intersections per incrementing radius (binned into 5 µm steps) was counted. For spine counts spine density was expressed as spines per 10 µm. Doublecortin cell counts are reported as unilateral estimates. Parametric tests (ANOVA and t-test) were used to compare groups. Greenhouse-Geisser adjustments were used to correct for sphericity for repeated measures analyses and Welch’s t-test for unequal variances were used where appropriate. One-sample t-tests comparing overall mean percentages against 50% were used to evaluate whether the lesion and sham groups were performing above chance on the T-maze task. SPSS software (version 20, IBM Corporation) was used to carry out statistical analyses.

Voxelwise analyses were performed in Matlab using an ANOVA script based on CoAxLab (2019). A two-by-four factorial ANOVA was implemented for FA and MD maps with a between factor of surgery (control /lesion) and the within factor of scan (one to four). Since false discovery rate correction eliminated virtually all voxels except for those in the lesion area. Data presented in **Fig. 6** are uncorrected results and restrict regional analyses to clusters below the α level of 0.005. Such identified voxels in the mammillothalamic tract, hippocampal, retrosplenial and mammillary body regions were subsequently used to elucidate the timeline of diffusivity changes across the four scans. A two-by-four ANOVA was performed for each region (SPSS 25.0, IBM, USA). In post-hoc analyses, both the differences from baseline (Scan 1) and between the groups (control/lesion) at each scan time were compared, and Bonferroni-adjusted. Unless otherwise stated, the threshold for significance was set at p < 0.05.

## Acknowledgments

This work was supported by the Wellcome Trust (090954/Z/09/Z; 102409/Z/13/A; 212273/Z/18/Z) and the Science Foundation Ireland (SFI 13/IA/2014). The authors thank Moira Davies, Aura Frizzati, Hadas Laufer, Tori Maddock, Andrew Nelson, Heather Phillips, Harry Potter, Andrew Stewart and Paul Wynne for technical, histological and surgical assistance.

## Supplementary Information

**Figure S1.**
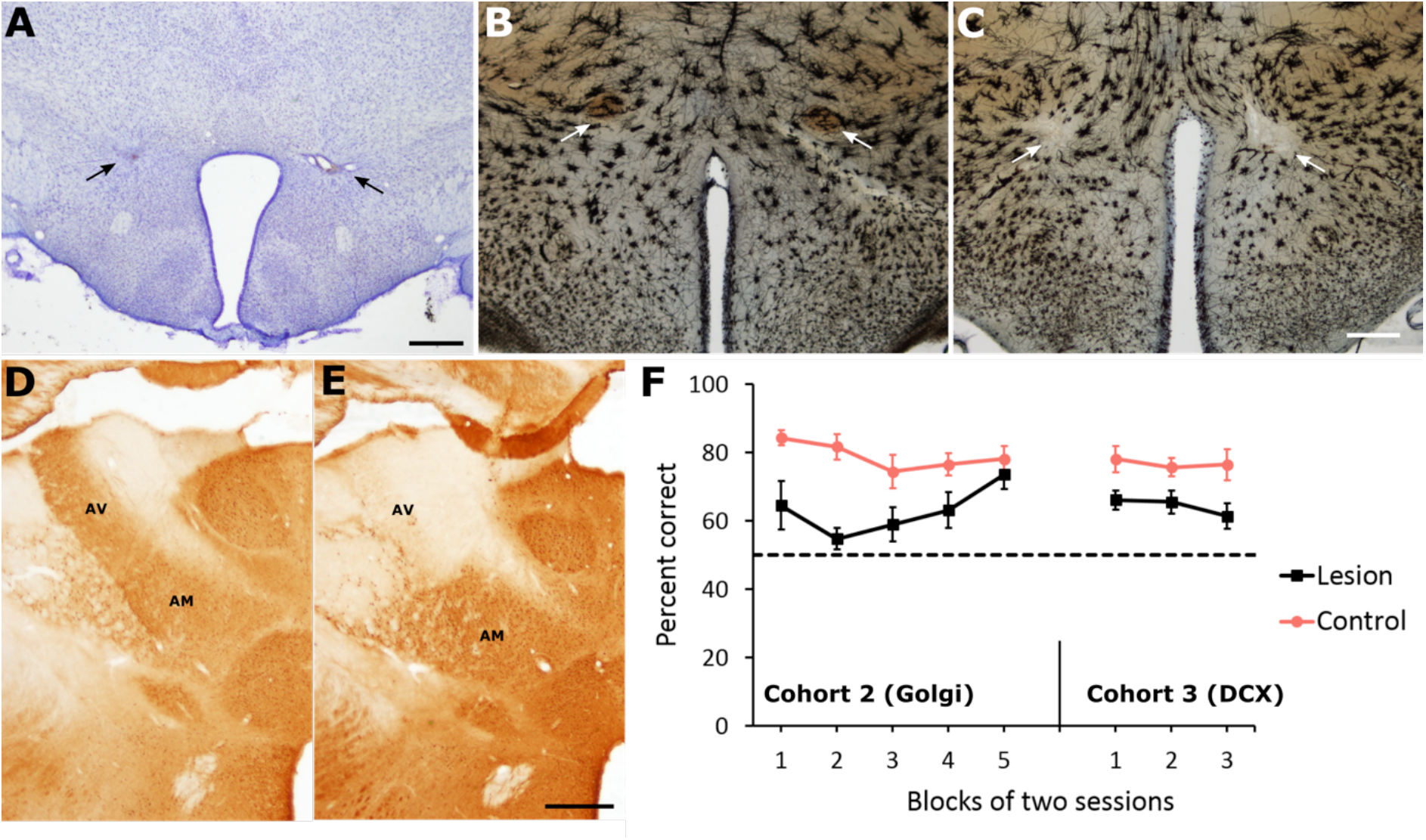
An additional representative mammillothalamic tract lesion from Experiment 2 (Cohort 3, **A**; Cohort 2, **B**, Control; **C**, Lesion). In Cohort 3, calbindin immunohistochemistry (**D**, Sham; **E**, Lesion) provided further validation of lesion success through an attenuation in staining in the anteroventral thalamic nucleus (AV). For technical reasons relating to the Golgi staining protocol it was not possible to perform calbindin immunohistochemistry in Cohort 2, however, behavioral performance on a reinforced alternation task in a T-maze was significantly reduced in both Cohort 2 (p<0.01; left facet of **F**) and cohort 3 (p<0.01; right facet of **F**). Scale bar = 500 µm.

**Figure S2.**
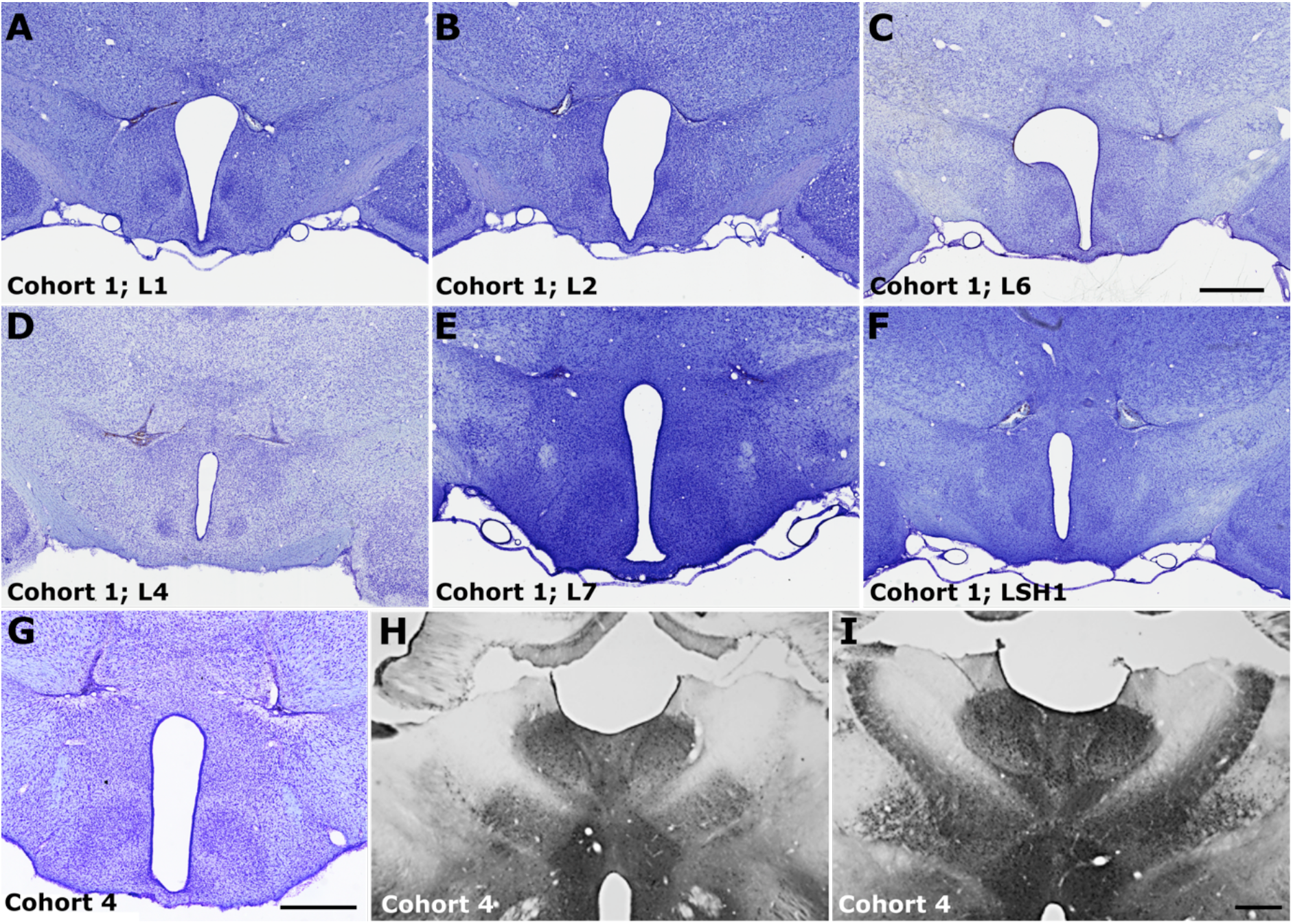
**A-F**) Nissl stained sections of mammillothalamic tract lesions from additional cases that contributed to Experiment 1 (Rat L5 shown in Fig. 1D). Rat identification labels refer to those shown in supplementary Fig. S3. **G**-**I**, An additional representative example of a mammillothalamic tract lesion (**G**), as well as low magnification photomicrographs showing the difference in calbindin immunoreactivity in the anteroventral thalamic nucleus of a lesion (**H**) and control (**I**) animal. Scale bar in A-F= 1 mm; G-I = 500 µm.

**Figure S3.**
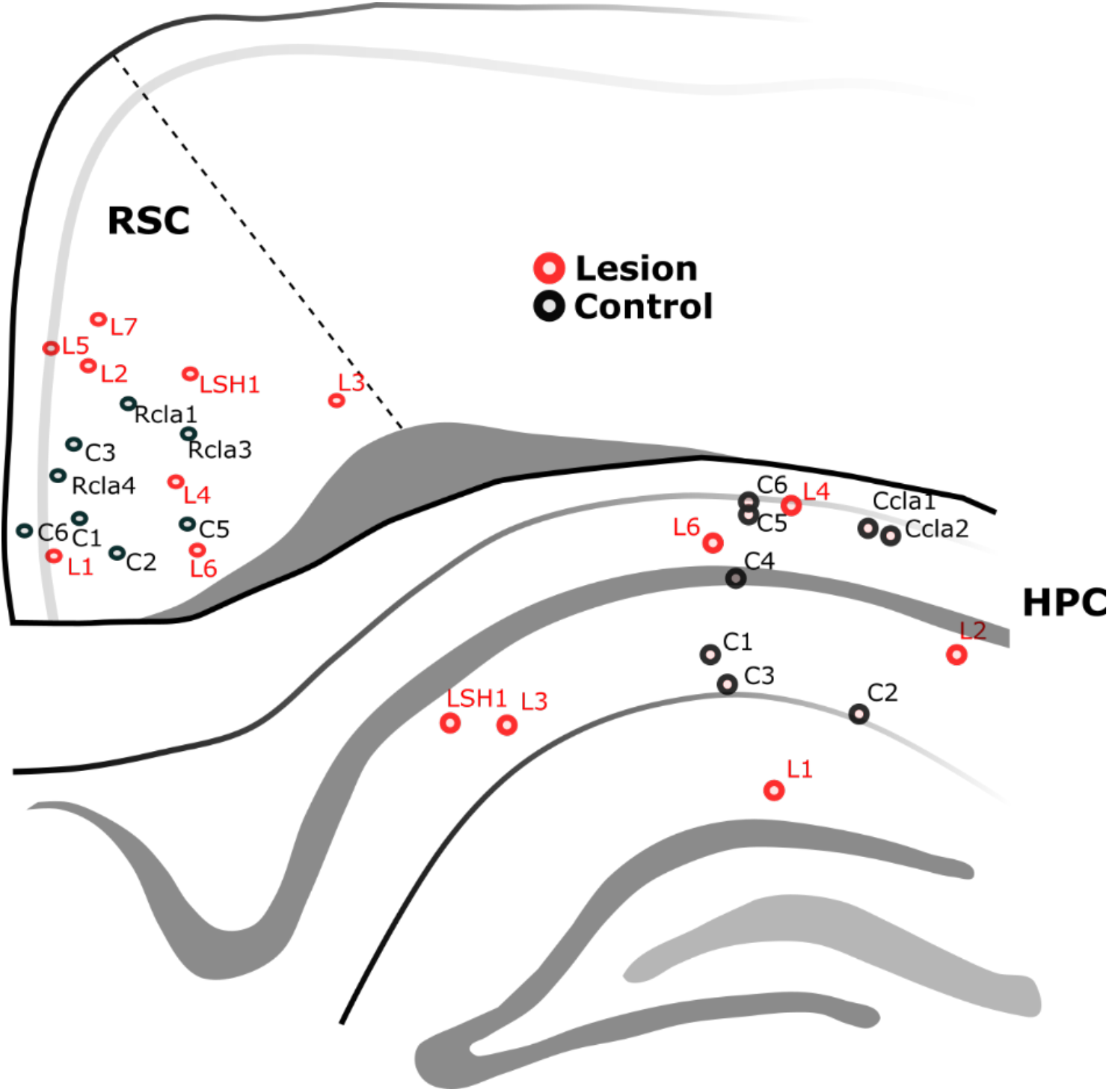
Schematic diagram showing the representative anterior-posterior position, and anatomical locations of all recording electrodes in the study (n= 30 in 18 animals). Local field potential recordings were made from the septal CA1 subregion of the hippocampus (HPC) and the septal retrosplenial cortex (RSC). Animal identification labels are provided to permit identification of the proportion of the animals that recorded from both regions simultaneously (n = 11), and those that were recorded from either region, singularly (n = 7).

**Figure S4.**
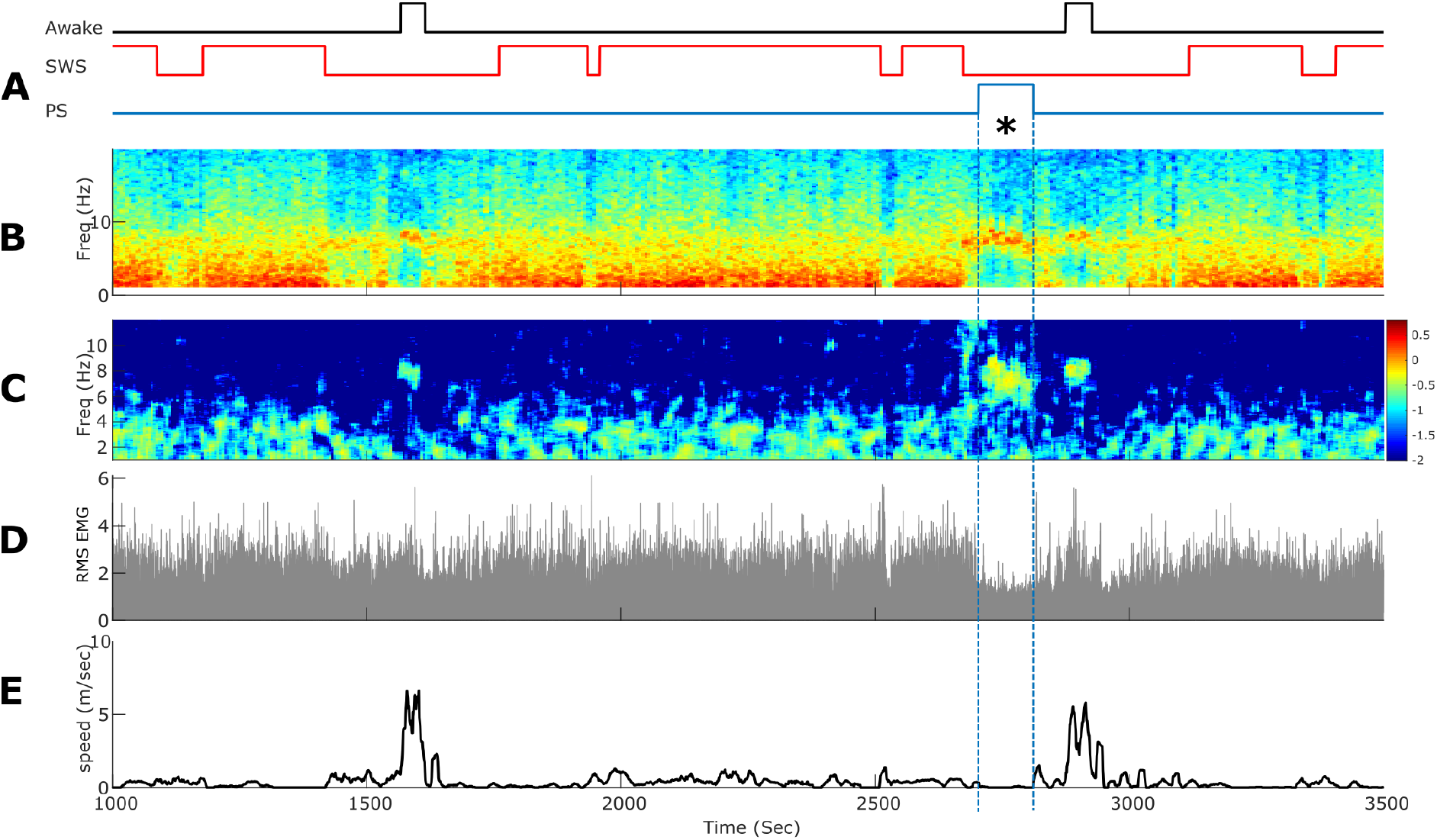
Episodes of paradoxical sleep were indexed (**A**; blue) from wakefulness, immobility (**A**, black) and slow-wave sleep (SWS; **A**, red) based on: 1. A threshold of a window-based (12 second with 50% overlap) theta-delta power ratio (**B**); 2. A threshold set to the root-mean square of the electromyography (EMG) signal (**D**), and 3. A speed threshold of 0 cm/second (**E**). **B** shows a representative recording session (from a control RSC electrode) dominated by slow-wave sleep, intermittent periods of wakefulness and an episode of PS. During PS (blue dashed line and asterisk), power in the theta band is dominant (**B**), running speed is zero **(E**) and EMG power is considerably reduced (**D**). C shows coherence in the time domain between local field potentials in RSC and HPC recorded simultaneously. During SWS, coherence is dominant in the delta band (1-4 Hz) and during wakefulness and PS, coherence is dominant in the theta band (4-12 Hz).

**Figure S5.**
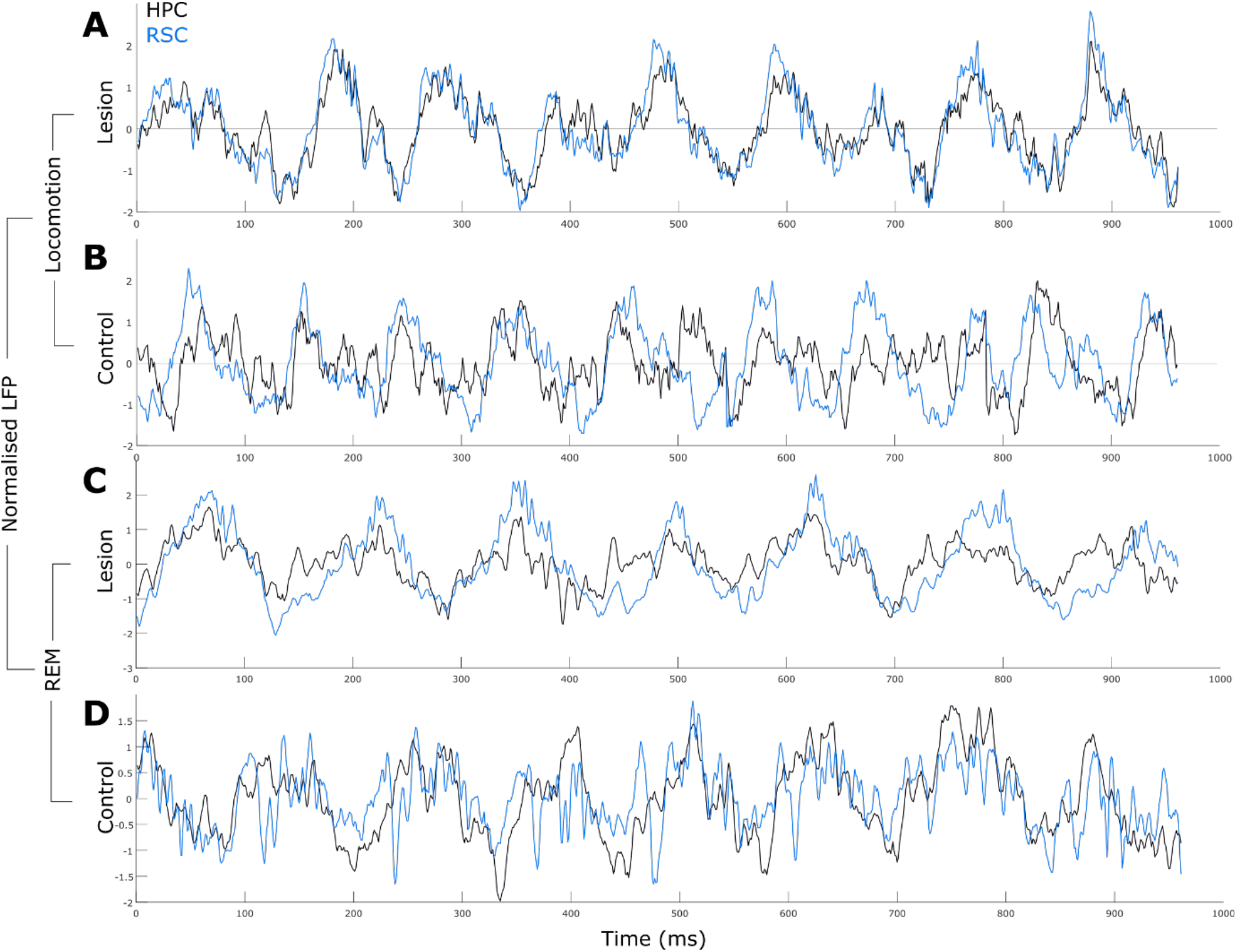
One second representative examples of local field potential traces recorded simultaneously from the retrosplenial cortex (RSC; blue) and the hippocampus (HPC; black). During active locomotion (**A**, **B**) and, paradoxical sleep (PS; **C**, **D**), animals with mammillothalamic tract lesions showed higher inter-regional coherence than controls.

**Figure S6.**
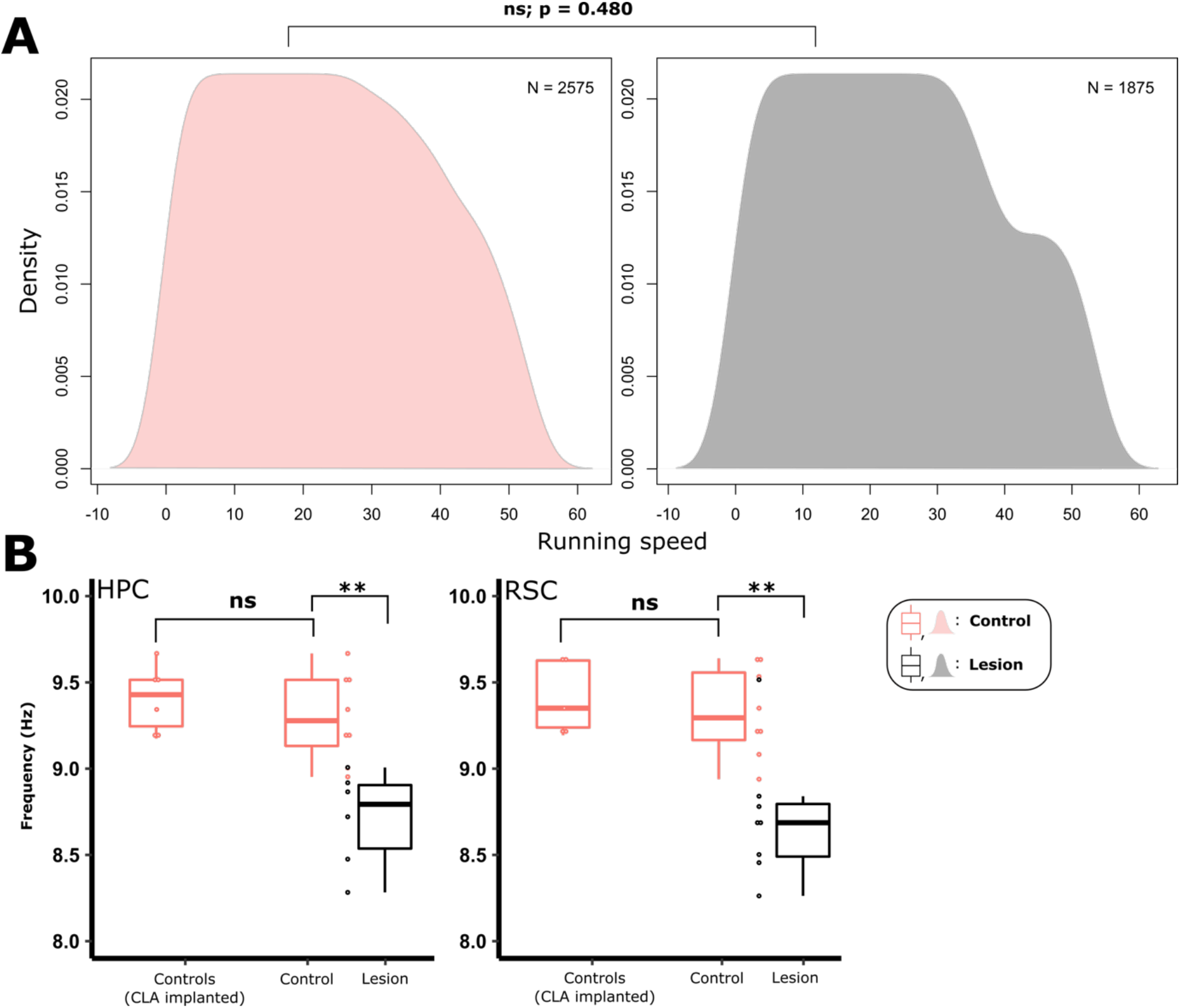
**A**, Kernel density distributions of speeds measured from control (red) and lesion (grey) groups between 0 and 55 cm/s. A permutation test of equality (Bowman and Azzalini, 1997) revealed no statistical difference in the distributions. **B**) Using the example of theta frequency during locomotion, control animals that were implanted in the claustrum as part of a separate project not augment the difference observed between control animals and lesion animals.

**Figure S7.**
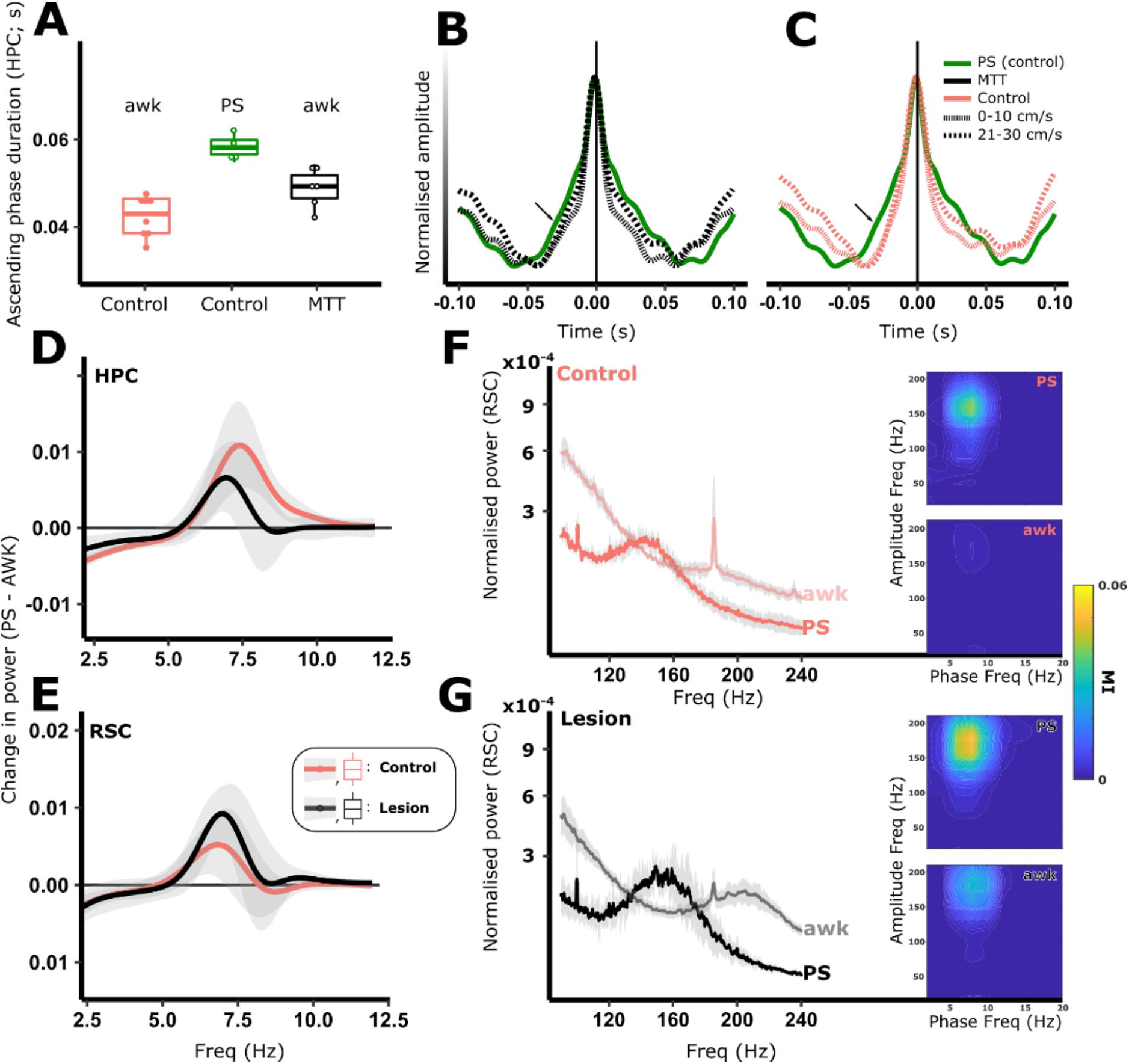
Both active locomotion and paradoxical sleep (PS) are ‘theta-rich’ states. Theta cycles in PS are more symmetrical (**A**-**C**) than during locomotion (AWK). In animals with lesions of the mammillothalamic tract (MTT), theta cycles during locomotion were closer in symmetry to PS cycles than those of controls (**A**-**C**). In the hippocampus (HPC), lesioned animals, the frequency of theta oscillations was reduced during both AWK and PS which is highlighted by the within-recording subtraction of the AWK power spectrum from that of PS (**D**). While theta frequency was reduced in RSC during AWK, it was not reduced during PS (**E**), however theta power was significantly increased in lesion animals (E; PS power spectrum minus AWK power spectrum). The observed increase in RSC power during PS was accompanied by an increase in the power and frequency of activity in the 100-180 Hz range (HFA; **F**-**G**), as well as an increase in phase amplitude coupling between theta and HFA (PS modulation index (MI) heatmaps in **F**-**G**). Increased phase-amplitude coupling present in the RSC during AWK (AWK MI heatmaps in **F**-**G**) was found to be due to a peak in HFA but at a considerably higher frequency (180-240 Hz) than that found during PS reinforcing the likelihood that distinct mechanisms, via independent neural populations, are responsible for the observed changes in awake locomotion and PS.

**Figure S8.**
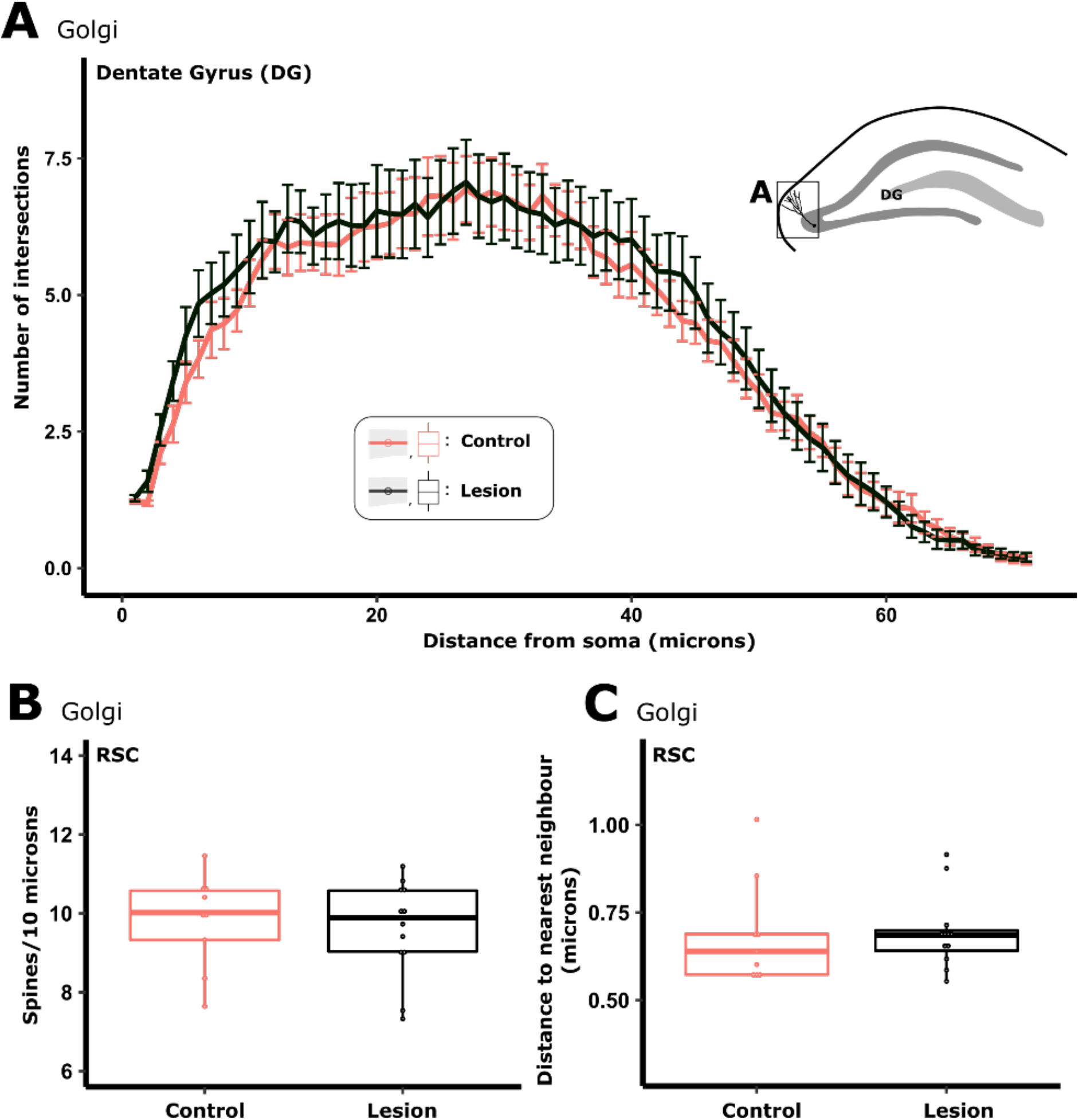
Sholl analysis of Golgi stained tissue revealed that the complexity of granule cells in the dentate gyrus of the hippocampal formation (DG) was not altered by lesions of the mammillothalamic tract (**A**). Analysis of spines on the apical dendrites of pyramidal cells in the retrosplenial granular cortex (RSC) showed that there was no difference in either the density, or the clustering of spines in animals with mammillothalamic tract lesions compared when compared with control animals.

**Figure S9.**
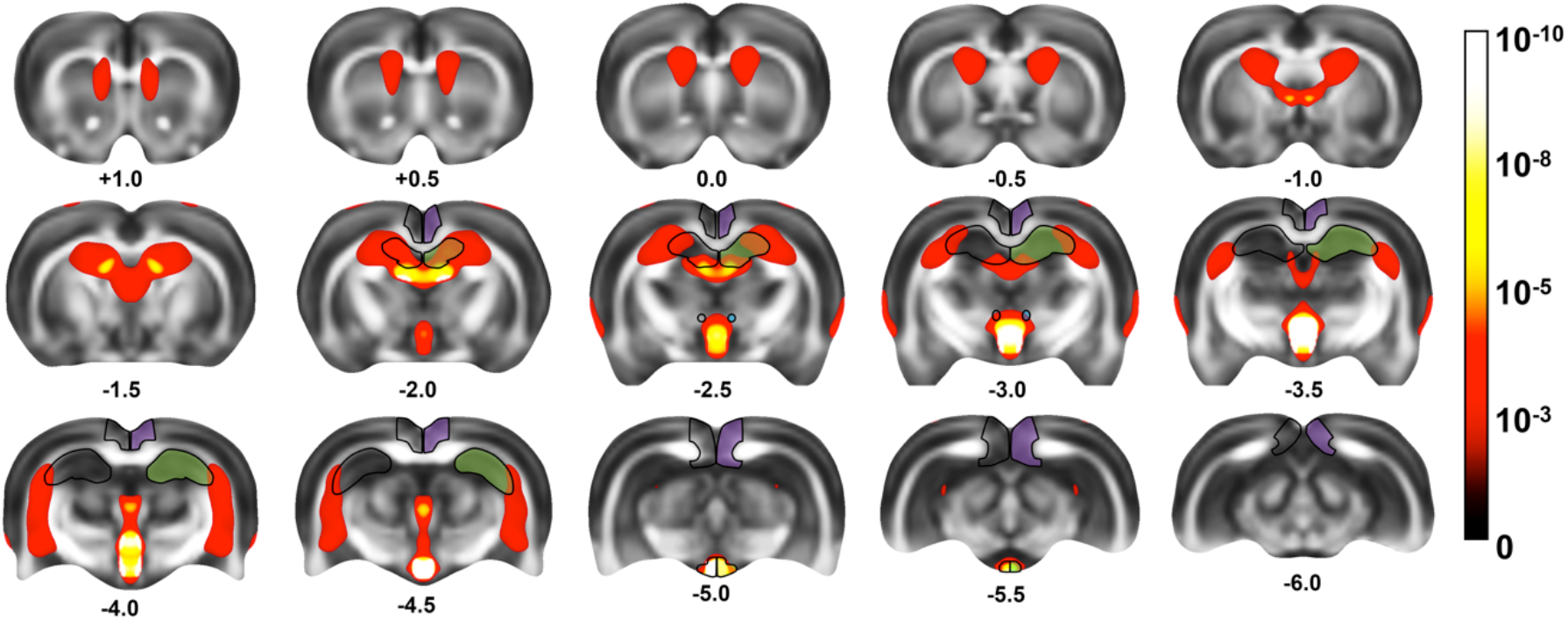
Changes in mean diffusivity (MD) elicited by MTT lesion and radial-arm maze training. The p-values (−Log_10_) represent the results of an ANOVA with the between factor of surgery (two levels) and the within factor of scan number (four levels). Significant interaction voxels have been thresholded to α < 0.01 and displayed using a heat map overlaid on top of a mean FA template. Successive coronal levels are numbered according to distance from bregma (mm).

**Figure S10.**
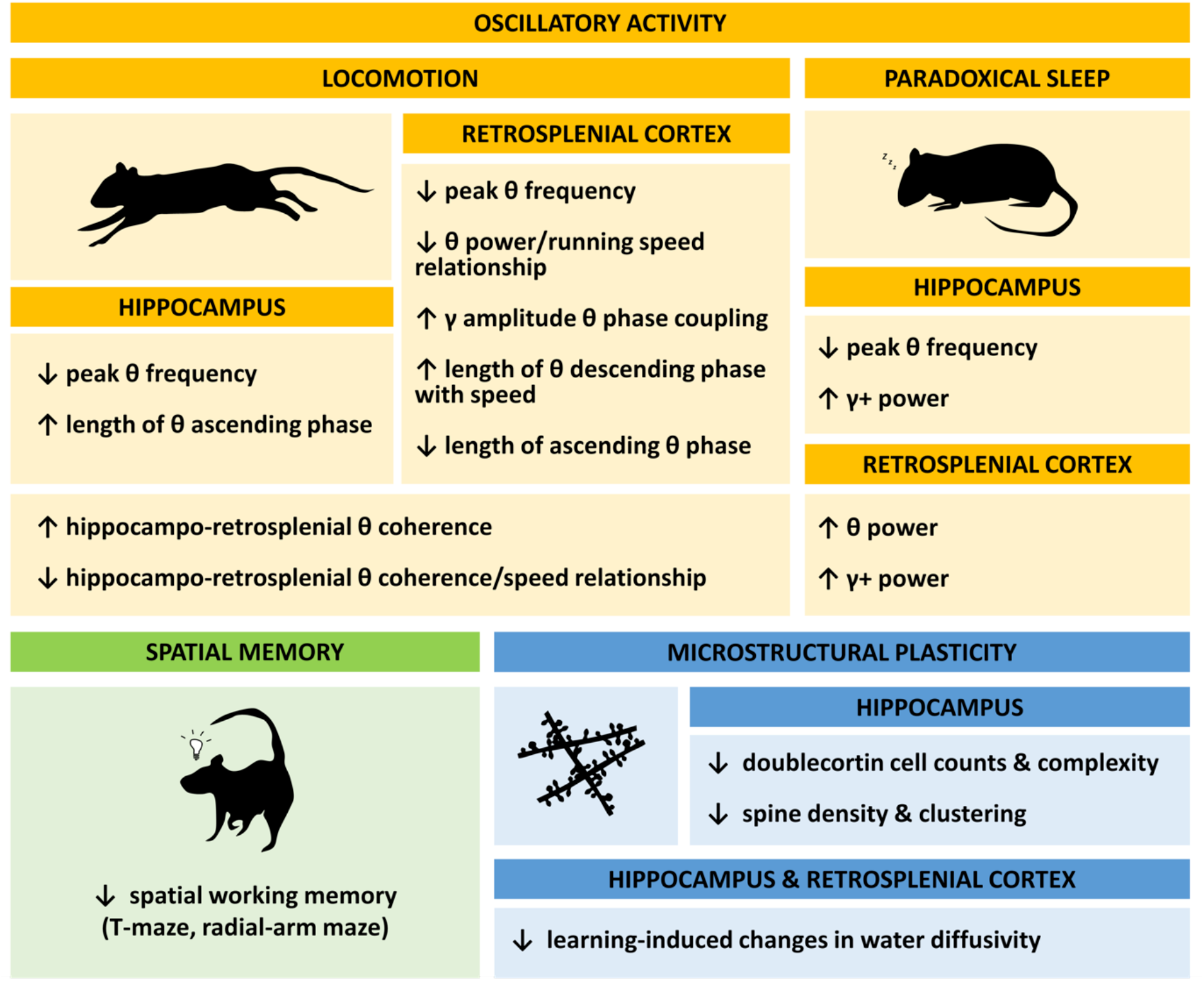
Summary of mammillothalamic tract lesion effects. Animals exhibited abnormal hippocampo-cortical oscillatory activity within theta and gamma bands: both within-region oscillatory activity and cross-regional synchrony were altered. The network changes were state-dependent, displaying different profiles during locomotion and paradoxical sleep. In addition, lesioned animals displayed microstructural changes, which appeared to reflect a suppression of learning-induced plasticity in lesioned animals.

**Table S1.** Precision of estimates for total DCX counts (unilateral)

|  | Sham | Lesion |
| --- | --- | --- |
| Mean | 12,626 | 8,742 |
| CE | 0.05 | 0.06 |
| CV | 0.18 | 0.14 |
| $CE^2 / CV^2$ | 0.08 | 0.17 |
| n | 11 | 8 |

## References

Aggleton JP, Brown MW. 1999. Episodic memory, amnesia, and the hippocampal-anterior thalamic axis. Behavioral and Brain Sciences 22:425–44; discussion 444-89.

Ahmed OJ, Mehta MR. 2012. Running speed alters the frequency of hippocampal gamma oscillations. J Neurosci 32:7373–83.

Avants BB, Epstein CL, Grossman M, Gee JC. 2008. Symmetric diffeomorphic image registration with cross-correlation: evaluating automated labeling of elderly and neurodegenerative brain. Med Image Anal 12:26–41.

Avants BB, Tustison NJ, Song G, Cook PA, Klein A, Gee JC. 2011. A reproducible evaluation of ANTs similarity metric performance in brain image registration. Neuroimage 54:2033–44.

Barbizet J. 1963. Defect of memorizing of hippocampal-mammillary origin: a review. J Neurol Neurosurg Psychiatry 26:127–35.

Belluscio MA, Mizuseki K, Schmidt R, Kempter R, Buzsaki G. 2012. Cross-frequency phase-phase coupling between theta and gamma oscillations in the hippocampus. J Neurosci 32:423–35.

Beracochea DJ, Micheau J, Jaffard R. 1995. Alteration of cortical and hippocampal cholinergic activities following lesion of the mammillary bodies in mice. Brain Res 670:53–8.

Bikbaev A, Manahan-Vaughan D. 2008. Relationship of hippocampal theta and gamma oscillations to potentiation of synaptic transmission. Front Neurosci 2:56–63.

Boyce R, Glasgow SD, Williams S, Adamantidis A. 2016. Causal evidence for the role of REM sleep theta rhythm in contextual memory consolidation. Science 352:812–6.

Briess D, Cotter D, Doshi R, Everall I. 1998. Mamillary body abnormalities in schizophrenia. Lancet 352:789–90.

Buzsaki G. 2002. Theta oscillations in the hippocampus. Neuron 33:325–40.

Buzsaki G, Rappelsberger P, Kellenyi L. 1985. Depth profiles of hippocampal rhythmic slow activity (’theta rhythm’) depend on behaviour. Electroencephalogr Clin Neurophysiol 61:77–88.

Cagnan H, Duff EP, Brown P. 2015. The relative phases of basal ganglia activities dynamically shape effective connectivity in Parkinson’s disease. Brain 138:1667–78.

Canolty RT, Knight RT. 2010. The functional role of cross-frequency coupling. Trends Cogn Sci 14:506–15.

Carlesimo GA, Lombardi MG, Caltagirone C. 2011. Vascular thalamic amnesia: a reappraisal. Neuropsychologia 49:777–789.

Carlesimo GA, Serra L, Fadda L, Cherubini A, Bozzali M, Caltagirone C. 2007. Bilateral damage to the mammillo-thalamic tract impairs recollection but not familiarity in the recognition process: a single case investigation. Neuropsychologia 45:2467–79.

Carpenter F, Burgess N, Barry C. 2017. Modulating medial septal cholinergic activity reduces medial entorhinal theta frequency without affecting speed or grid coding. Sci Rep 7:14573.

Chang LC, Jones DK, Pierpaoli C. 2005. RESTORE: robust estimation of tensors by outlier rejection. Magn Reson Med 53:1088–95.

Clarke S, Assal G, Bogousslavsky J, Regli F, Townsend DW, Leenders KL, Blecic S. 1994. Pure amnesia after unilateral left polar thalamic infarct: topographic and sequential neuropsychological and metabolic (PET) correlations. Journal of Neurology, Neurosurgery & Psychiatry 57:27–34.

Cole SR, Voytek B. 2018. Hippocampal theta bursting and waveform shape reflect CA1 spiking patterns. bioRxiv:452987.

Couillard-Despres S, Winner B, Schaubeck S, Aigner R, Vroemen M, Weidner N, Bogdahn U, Winkler J, Kuhn HG, Aigner L. 2005. Doublecortin expression levels in adult brain reflect neurogenesis. Eur J Neurosci 21:1–14.

Delay J, Brion S. 1969. Le Syndrome de Korsakoff. Paris: Masson.

Denby CE, Vann SD, Tsivilis D, Aggleton JP, Montaldi D, Roberts N, Mayes AR. 2009. The frequency and extent of mammillary body atrophy associated with surgical removal of a colloid cyst. AJNR Am J Neuroradiol 30:736–43.

Dillingham CM, Frizzati A, Nelson AJ, Vann SD. 2015. How do mammillary body inputs contribute to anterior thalamic function? Neurosci Biobehav Rev 54:108–19.

Dorph-Petersen KA, Nyengaard JR, Gundersen HJ. 2001. Tissue shrinkage and unbiased stereological estimation of particle number and size. J Microsc 204:232–46.

Dudchenko PA. 2001. How do animals actually solve the T maze? Behav Neurosci 115:850–60.

Dupret D, Revest JM, Koehl M, Ichas F, De Giorgi F, Costet P, Abrous DN, Piazza PV. 2008. Spatial relational memory requires hippocampal adult neurogenesis. PLoS One 3:e1959.

Dzieciol AM, Bachevalier J, Saleem KS, Gadian DG, Saunders R, Chong WKK, Banks T, Mishkin M, Vargha-Khadem F. 2017. Hippocampal and diencephalic pathology in developmental amnesia. Cortex 86:33–44.

Farioli-Vecchioli S, Saraulli D, Costanzi M, Pacioni S, Cina I, Aceti M, Micheli L, Bacci A, Cestari V, Tirone F. 2008. The timing of differentiation of adult hippocampal neurons is crucial for spatial memory. PLoS Biol 6:e246.

Feldman HM, Yeatman JD, Lee ES, Barde LH, Gaman-Bean S. 2010. Diffusion tensor imaging: a review for pediatric researchers and clinicians. J Dev Behav Pediatr 31:346–56.

Fuhrmann F, Justus D, Sosulina L, Kaneko H, Beutel T, Friedrichs D, Schoch S, Schwarz MK, Fuhrmann M, Remy S. 2015. Locomotion, Theta Oscillations, and the Speed-Correlated Firing of Hippocampal Neurons Are Controlled by a Medial Septal Glutamatergic Circuit. Neuron 86:1253–64.

Gamper E. 1928. Zur Frage der Polioencephalitis der chronischen Alkoholiker. Anatomische Befunde beim chronischem Korsakow und ihre Beziehungen zum klinischen Bild. Deutsche Z. Nervenheilkd 102:122–129.

Gardini S, Venneri A, Sambataro F, Cuetos F, Fasano F, Marchi M, Crisi G, Caffarra P. 2015. Increased functional connectivity in the default mode network in mild cognitive impairment: a maladaptive compensatory mechanism associated with poor semantic memory performance. J Alzheimers Dis 45:457–70.

Gonzalo-Ruiz A, Morte L. 2000. Localization of amino acids, neuropeptides and cholinergic markers in neurons of the septum-diagonal band complex projecting to the retrosplenial granular cortex of the rat. Brain Res Bull 52:499–510.

Goyal A, Miller J, Qasim S, Watrous AJ, Stein JM, Inman CS, Gross RE, Willie JT, Lega B, Lin J-J and others. 2018. Functionally distinct high and low theta oscillations in the human hippocampus. bioRxiv:498055.

Gudden H. 1896. Klinische und anatomische Beitrage zur Kenntnis der multiplen Alkoholneuritis nebst Bernerkungen uber die Regenerationsvorgange im peripheren Nervensystem. Arch Psychiatr 28:643–741.

Gundersen HJ. 1986. Stereology of arbitrary particles. A review of unbiased number and size estimators and the presentation of some new ones, in memory of William R. Thompson. J Microsc 143:3–45.

Gundersen HJ, Jensen EB, Kieu K, Nielsen J. 1999. The efficiency of systematic sampling in stereology--reconsidered. J Microsc 193:199–211.

Harland BC, Collings DA, McNaughton N, Abraham WC, Dalrymple-Alford JC. 2014. Anterior thalamic lesions reduce spine density in both hippocampal CA1 and retrosplenial cortex, but enrichment rescues CA1 spines only. Hippocampus 24:1232–47.

Harris KD, Thiele A. 2011. Cortical state and attention. Nat Rev Neurosci 12:509–23.

Hawellek DJ, Hipp JF, Lewis CM, Corbetta M, Engel AK. 2011. Increased functional connectivity indicates the severity of cognitive impairment in multiple sclerosis. Proc Natl Acad Sci U S A 108:19066–71.

Huerta PT, Lisman JE. 1993. Heightened synaptic plasticity of hippocampal CA1 neurons during a cholinergically induced rhythmic state. Nature 364:723–5.

Jankowski MM, Ronnqvist KC, Tsanov M, Vann SD, Wright NF, Erichsen JT, Aggleton JP, O’Mara SM. 2013. The anterior thalamus provides a subcortical circuit supporting memory and spatial navigation. Front Syst Neurosci 7:45.

Jeewajee A, Lever C, Burton S, O’Keefe J, Burgess N. 2008. Environmental novelty is signaled by reduction of the hippocampal theta frequency. Hippocampus 18:340–8.

Jones DK, Horsfield MA, Simmons A. 1999. Optimal strategies for measuring diffusion in anisotropic systems by magnetic resonance imaging. Magn Reson Med 42:515–25.

Kastellakis G, Cai DJ, Mednick SC, Silva AJ, Poirazi P. 2015. Synaptic clustering within dendrites: an emerging theory of memory formation. Prog Neurobiol 126:19–35.

Kempermann G, Jessberger S, Steiner B, Kronenberg G. 2004. Milestones of neuronal development in the adult hippocampus. Trends Neurosci 27:447–52.

Kikinis Z, Makris N, Sydnor VJ, Bouix S, Pasternak O, Coman IL, Antshel KM, Fremont W, Kubicki MR, Shenton ME and others. 2019. Abnormalities in gray matter microstructure in young adults with 22q11.2 deletion syndrome. Neuroimage Clin 21:101611.

Kirk IJ. 1998. Frequency modulation of hippocampal theta by the supramammillary nucleus, and other hypothalamo-hippocampal interactions: mechanisms and functional implications. Neurosci Biobehav Rev 22:291–302.

Kocsis B, Vertes RP. 1994. Characterization of neurons of the supramammillary nucleus and mammillary body that discharge rhythmically with the hippocampal theta rhythm in the rat. J Neurosci 14:7040–52.

Koike BDV, Farias KS, Billwiller F, Almeida-Filho D, Libourel PA, Tiran-Cappello A, Parmentier R, Blanco W, Ribeiro S, Luppi PH and others. 2017. Electrophysiological Evidence That the Retrosplenial Cortex Displays a Strong and Specific Activation Phased with Hippocampal Theta during Paradoxical (REM) Sleep. J Neurosci 37:8003–8013.

Kopelman MD. 1995. The Korsakoff syndrome. The British Journal of Psychiatry 166:154–173.

Kumar R, Birrer BVX, Macey PM, Woo MA, Gupta RK, Yan-Go FL, Harper RM. 2008. Reduced mammillary body volume in patients with obstructive sleep apnea. Neurosci Lett 438:330–334.

Kumar R, Woo MA, Birrer BVX, Macey PM, Fonarow GC, Hamilton MA, Harper RM. 2009. Mammillary bodies and fornix fibers are injured in heart failure. Neurobiology of Disease 33:236–242.

Lee H, Fell J, Axmacher N. 2013. Electrical engram: how deep brain stimulation affects memory. Trends Cogn Sci 17:574–84.

Leemans A, B J, J S, Jones DK. 2009. ExploreDTI: a graphical toolbox for processing, analyzing, and visualizing diffusion MR data. 17th Annual Meeting of Int Soc Magn Reson Med Hawaii, USA:3537.

Leemans A, Jones DK. 2009. The B-matrix must be rotated when correcting for subject motion in DTI data. Magn Reson Med 61:1336–49.

Lemaire V, Tronel S, Montaron MF, Fabre A, Dugast E, Abrous DN. 2012. Long-lasting plasticity of hippocampal adult-born neurons. J Neurosci 32:3101–8.

Leung LW, Lopes da Silva FH, Wadman WJ. 1982. Spectral characteristics of the hippocampal EEG in the freely moving rat. Electroencephalogr Clin Neurophysiol 54:203–19.

Longair MH, Baker DA, Armstrong JD. 2011. Simple Neurite Tracer: open source software for reconstruction, visualization and analysis of neuronal processes. Bioinformatics 27:2453–4.

Mandyam CD, Wee S, Crawford EF, Eisch AJ, Richardson HN, Koob GF. 2008. Varied access to intravenous methamphetamine self-administration differentially alters adult hippocampal neurogenesis. Biol Psychiatry 64:958–65.

Maurer AP, Vanrhoads SR, Sutherland GR, Lipa P, McNaughton BL. 2005. Self-motion and the origin of differential spatial scaling along the septo-temporal axis of the hippocampus. Hippocampus 15:841–52.

Merker B. 2013. Cortical gamma oscillations: the functional key is activation, not cognition. Neurosci Biobehav Rev 37:401–17.

Milczarek MM, Vann SD, Sengpiel F. 2018. Spatial Memory Engram in the Mouse Retrosplenial Cortex. Curr Biol 28:1975–1980 e6.

Mitra P, Bokil H. 2008. Observed brain dynamics. New York; Oxford: Oxford University Press.

Moser MB, Trommald M, Andersen P. 1994. An increase in dendritic spine density on hippocampal CA1 pyramidal cells following spatial learning in adult rats suggests the formation of new synapses. Proc Natl Acad Sci U S A 91:12673–5.

Moser MB, Trommald M, Egeland T, Andersen P. 1997. Spatial training in a complex environment and isolation alter the spine distribution differently in rat CA1 pyramidal cells. J Comp Neurol 380:373–81.

Nelson AJ, Vann SD. 2014. Mammilliothalamic tract lesions disrupt tests of visuo-spatial memory. Behav Neurosci 128:494–503.

Nokia MS, Sisti HM, Choksi MR, Shors TJ. 2012. Learning to learn: theta oscillations predict new learning, which enhances related learning and neurogenesis. PLoS One 7:e31375.

Olvera-Cortes E, Cervantes M, Gonzalez-Burgos I. 2002. Place-learning, but not cue-learning training, modifies the hippocampal theta rhythm in rats. Brain Res Bull 58:261–70.

Orr G, Rao G, Houston FP, McNaughton BL, Barnes CA. 2001. Hippocampal synaptic plasticity is modulated by theta rhythm in the fascia dentata of adult and aged freely behaving rats. Hippocampus 11:647–54.

Osher S, Shen J. 2000. Digitized PDE Method for Data Restoration. In: Anastassiou G, editor. Analytic-Computational Methods in Applied Mathematics. p 751–771.

Ozyurt J, Thiel CM, Lorenzen A, Gebhardt U, Calaminus G, Warmuth-Metz M, Muller HL. 2014. Neuropsychological outcome in patients with childhood craniopharyngioma and hypothalamic involvement. J Pediatr 164:876–881 e4.

Pan WX, McNaughton N. 1997. The medial supramammillary nucleus, spatial learning and the frequency of hippocampal theta activity. Brain Res 764:101–8.

Papez JW. 1937. A proposed mechanism of emotion. Arch. Neurol. Psychiatry 38:725–743.

Paxinos G, Watson C. 1998. The rat brain in stereotaxic coordinates. San Diego; London: Academic.

Perry JC, Pakkenberg B, Vann SD. 2019. Striking reduction in neurons and glial cells in anterior thalamic nuclei of older patients with Down syndrome. Neurobiol Aging 75:54–61.

Pierpaoli C, Basser PJ. 1996. Toward a quantitative assessment of diffusion anisotropy. Magn Reson Med 36:893–906.

Ray S, Maunsell JH. 2011. Different origins of gamma rhythm and high-gamma activity in macaque visual cortex. PLoS Biol 9:e1000610.

Renouard L, Billwiller F, Ogawa K, Clement O, Camargo N, Abdelkarim M, Gay N, Scote-Blachon C, Toure R, Libourel PA and others. 2015. The supramammillary nucleus and the claustrum activate the cortex during REM sleep. Sci Adv 1:e1400177.

Richard GR, Titiz A, Tyler A, Holmes GL, Scott RC, Lenck-Santini PP. 2013. Speed modulation of hippocampal theta frequency correlates with spatial memory performance. Hippocampus 23:1269–79.

Rogerson T, Cai DJ, Frank A, Sano Y, Shobe J, Lopez-Aranda MF, Silva AJ. 2014. Synaptic tagging during memory allocation. Nat Rev Neurosci 15:157–69.

Sagi Y, Tavor I, Hofstetter S, Tzur-Moryosef S, Blumenfeld-Katzir T, Assaf Y. 2012. Learning in the fast lane: new insights into neuroplasticity. Neuron 73:1195–203.

Sarra S. 2006. Digital total variation filtering as postprocessing for Chebyshev pseudospectral methods for conservation laws.. Numerical Algorithms 41:17–33.

Savastano LE, Hollon TC, Barkan AL, Sullivan SE. 2018. Korsakoff syndrome from retrochiasmatic suprasellar lesions: rapid reversal after relief of cerebral compression in 4 cases. J Neurosurg 128:1731–1736.

Scheffer-Teixeira R, Belchior H, Leao RN, Ribeiro S, Tort AB. 2013. On high-frequency field oscillations (>100 Hz) and the spectral leakage of spiking activity. J Neurosci 33:1535–9.

Schindelin J, Arganda-Carreras I, Frise E, Kaynig V, Longair M, Pietzsch T, Preibisch S, Rueden C, Saalfeld S, Schmid B and others. 2012. Fiji: an open-source platform for biological-image analysis. Nat Methods 9:676–82.

Sharp PE, Koester K. 2008. Lesions of the mammillary body region alter hippocampal movement signals and theta frequency: implications for path integration models. Hippocampus 18:862–78.

Sheremet A, Burke SN, Maurer AP. 2016. Movement Enhances the Nonlinearity of Hippocampal Theta. J Neurosci 36:4218–30.

Sheremet A, Kennedy JP, Qin Y, Zhou Y, Lovett SD, Burke SN, Maurer AP. 2019. Theta-gamma cascades and running speed. J Neurophysiol 121:444–458.

Slawinska U, Kasicki S. 1998. The frequency of rat’s hippocampal theta rhythm is related to the speed of locomotion. Brain Res 796:327–31.

Sorzano CO, Thevenaz P, Unser M. 2005. Elastic registration of biological images using vector-spline regularization. IEEE Trans Biomed Eng 52:652–63.

Talk A, Kang E, Gabriel M. 2004. Independent generation of theta rhythm in the hippocampus and posterior cingulate cortex. Brain Res 1015:15–24.

Tavor I, Hofstetter S, Assaf Y. 2013. Micro-structural assessment of short term plasticity dynamics. Neuroimage 81:1–7.

Tort AB, Komorowski R, Eichenbaum H, Kopell N. 2010. Measuring phase-amplitude coupling between neuronal oscillations of different frequencies. J Neurophysiol 104:1195–210.

Tronel S, Fabre A, Charrier V, Oliet SH, Gage FH, Abrous DN. 2010. Spatial learning sculpts the dendritic arbor of adult-born hippocampal neurons. Proc Natl Acad Sci U S A 107:7963–8.

Tsanov M, Manahan-Vaughan D. 2009. Long-term plasticity is proportional to theta-activity. PLoS One 4:e5850.

Tsanov M, Wright N, Vann SD, Erichsen JT, Aggleton JP, O’Mara SM. 2011. Hippocampal inputs mediate theta-related plasticity in anterior thalamus. Neuroscience 187:52–62.

Van der Werf YD, Scheltens P, Lindeboom J, Witter MP, Uylings HBM, Jolles J. 2003. Deficits of memory, executive functioning and attention following infarction in the thalamus; a study of 22 cases with localised lesions. Neuropsychologia 41:1330–1344.

Van der Werf YD, Witter MP, Uylings HB, Jolles J. 2000. Neuropsychology of infarctions in the thalamus: a review. Neuropsychologia 38:613–27.

Vann SD. 2009. Gudden’s ventral tegmental nucleus is vital for memory: re-evaluating diencephalic inputs for amnesia. Brain 132:2372–84.

Vann SD. 2010. Re-evaluating the role of the mammillary bodies in memory. Neuropsychologia 48:2316–27.

Vann SD. 2013. Dismantling the Papez circuit for memory in rats. Elife 2:e00736.

Vann SD, Aggleton JP. 2003. Evidence of a spatial encoding deficit in rats with lesions of the mammillary bodies or mammillothalamic tract. J Neurosci 23:3506–14.

Vann SD, Albasser MM. 2009. Hippocampal, retrosplenial, and prefrontal hypoactivity in a model of diencephalic amnesia: Evidence towards an interdependent subcortical-cortical memory network. Hippocampus 19:1090–102.

Vann SD, Nelson AJ. 2015. The mammillary bodies and memory: more than a hippocampal relay. Prog Brain Res 219:163–85.

Vann SD, Tsivilis D, Denby CE, Quamme JR, Yonelinas AP, Aggleton JP, Montaldi D, Mayes AR. 2009. Impaired recollection but spared familiarity in patients with extended hippocampal system damage revealed by 3 convergent methods. Proceedings of the Academy of Natural Sciences 106:5442–7.

Vukovic J, Borlikova GG, Ruitenberg MJ, Robinson GJ, Sullivan RK, Walker TL, Bartlett PF. 2013. Immature doublecortin-positive hippocampal neurons are important for learning but not for remembering. J Neurosci 33:6603–13.

West MJ, Slomianka L, Gundersen HJ. 1991. Unbiased stereological estimation of the total number of neurons in thesubdivisions of the rat hippocampus using the optical fractionator. Anat Rec 231:482–97.

Wolff M, Vann SD. 2019. The Cognitive Thalamus as a Gateway to Mental Representations. J Neurosci 39:3–14.

Yoneoka Y, Takeda N, Inoue A, Ibuchi Y, Kumagai T, Sugai T, Takeda K, Ueda K. 2004. Acute Korsakoff syndrome following mammillothalamic tract infarction. AJNR Am J Neuroradiol 25:964–8.

Zakowski W, Zawistowski P, Braszka L, Jurkowlaniec E. 2017. The effect of pharmacological inactivation of the mammillary body and anterior thalamic nuclei on hippocampal theta rhythm in urethane-anesthetized rats. Neuroscience 362:196–205.

## Supplementary References

Bowman, A.W. and Azzalini, A. (1997). Applied Smoothing Techniques for Data Analysis: the Kernel Approach with S-Plus Illustrations. Oxford University Press, Oxford.

